# Infection of 5xFAD mice with a mouse-adapted SARS-CoV-2 does not alter Alzheimer’s disease neuropathology yet induces wide-spread changes in gene expression across diverse cell types

**DOI:** 10.64898/2025.12.19.695600

**Authors:** Susana Furman, Latifa Zayou, Kate Inman Tsourmas, Dominic Ibarra Javonillo, Gema M. Olivarria, Yuting Cheng, Collin Pachow, Kellie Fernandez, Lucas Le, Robert A. Edwards, Dequina A. Nicholas, Gabriela Pacheco Sanchez, Ralph S. Baric, Kim N. Green, Thomas E. Lane

**Author notes:** <u>Address Correspondence:</u> Thomas E. Lane, Ph.D. 2213 McGaugh Hall, University of California, Irvine Irvine, CA 92697-4545, USA. <u>Funding Information</u> Alzheimer’s Association grant 105190 (TEL) National Institutes of Health (NIH-NINDS) grant R35NS116835 (TEL) National Institutes of Health (NIH-NIA) grant R01AG081599 (KNG) National Institutes of Health (NIH-NIA) grant U54 AG054349 (KNG) National Institutes of Health (NIH-NIAID) R01AI110700 (RSB) National Cancer Institute (NCI) 5P30CA062203-26 (RAE) National Institutes of Health (NIH-NIAID) U01AI180164 (DAN) National Institutes of Health (NIH-NINDS) Training Grant T32 NS121727 (DIJ) National Institutes of Health (NIH-AI) Training Grant 5T32 AI007319-33 (GMO).

## Abstract

Alzheimer’s disease (AD) is a progressive neurodegenerative disorder and the leading cause of dementia. It is characterized by cognitive decline and accumulation of amyloid beta (Aβ) plaques and neurofibrillary tangles. Accumulating evidence indicates that viral infection may worsen and/or increase development of established AD pathology. The COVID-19 pandemic has brought attention to the link between SARS-CoV-2 infection and neurologic conditions that vary in severity and duration, as well as the worsening of clinical symptoms in elderly people with dementia. To better understand potential mechanisms by which SARS-CoV-2 infection impacts AD neuropathology, aged 5xFAD and wildtype (WT) mice were intranasally infected with mouse-adapted SARS-CoV-2 (MA10). Intranasal infection of aged-matched (10-14 month) 5xFAD or wild type (WT) C57BL/6 mice with MA10 resulted in viral infection of the lungs that correlated with acute viral pneumonia characterized by lymphocyte inflammation and antiviral immune responses. Viral RNA was not detected within the central nervous system (CNS) of either WT or 5xFAD mice at days 7 or 21 post-infection (p.i.), nor were there signs of overt glial activation or neuroinflammation. There were no differences in either Aβ plaque volume or number within the brains of MA10-infected 5xFAD mice compared to uninfected 5xFAD mice. However, bulk RNA sequencing and spatial transcriptomics revealed evidence of altered expression of genes associated with neuronal and glial dysfunction, as well as reduced expression of genes encoding adhesion molecules in vascular endothelial cells. Collectively, these findings demonstrate that MA10 infection did not affect Aβ plaque size or numbers in 5xFAD mice, yet in both WT and 5xFAD mice, there were numerous down-stream effects on gene expression associated with resident CNS cell function.

## Introduction

Alzheimer’s disease (AD) is the most common cause of age-related dementia and impairs memory formation and other intellectual functions, leading to a decline in quality of life and ability to perform daily tasks. Pathology includes the aberrant accumulation of amyloid plaques and neurofibrillary tangles, accompanied with neuronal loss and neuroinflammation ^1^. AD is a complex multifactorial disease, where a wide range of risk factors are likely to influence its onset. While age has been established as the most important risk factor ^2^, a wide variety of genetic ^3–7^ and environmental factors ^8–14^ have been associated with an increased risk of AD. Some studies have suggested possible links between viral infection and acceleration of onset or exacerbation of established pathology and clinical symptoms associated with various neurodegenerative diseases, including AD^15,16^. Microbial pathogens have been suggested to contribute to AD pathogenesis, and numerous studies have implicated several infectious agents in AD^17–22^. Viruses and viral proteins may have amyloidogenic properties that can lead to the formation of amyloid aggregates^23–25^. The increasing number of studies associating infections to neurodegeneration and AD emphasizes the importance of continued investigation on the contribution of microbial infection to AD pathology and other dementias.

COVID-19 is caused by the severe acute respiratory syndrome coronavirus 2 (SARS-CoV-2). In 2020, COVID-19 rapidly developed into a global pandemic ^26^ that affected hundreds of millions of people worldwide ^27^. In the majority of individuals, infection presents with mild-to-moderate respiratory disease and other flu-like symptoms^28,29^, with more severe presentations resulting in acute respiratory distress syndrome (ARDS) that could result in respiratory failure ^30^. Although COVID-19 symptoms are mostly respiratory in nature, neurological symptoms and complications have also been documented ^31–35^. Common neurological symptoms include headache, dizziness, as well as olfactory and gustatory dysfunction^36–38^. However, COVID-19 patients with more severe disease have also been associated with ischemic strokes, cerebrovascular disease, coagulation abnormalities, seizures, meningitis and encephalitis^34,39–46^.

Several risk factors have been associated with an increase in COVID-19 clinical disease, and evidence suggests that increased age is associated with not only a higher risk of disease severity, but also mortality^47–49^. Pre-existing dementia also appears to be a contributing risk factor for COVID-19. Infection is associated with a worsening of clinical symptoms in elderly patients with dementia and increased mortality^50,51^, and among patients with dementia, AD was the most prevalent cognitive impairment in those who died of COVID-19 ^52^. COVID-19 appears to impact brain structure and function, and while definitive mechanisms between COVID-19 and neurodegeneration remain poorly understood, several studies suggest that COVID-19 may induce or accelerate neurodegenerative processes, including those related to dementia and AD ^53^. Recent evidence suggests that various viruses, including coronaviruses like SARS-CoV-1 ^24^ and more recently SARS-CoV-2 ^54^ possess viral proteins that exhibit amyloidogenic properties and can lead to the formation of amyloid aggregates. In addition, infection may alter resident CNS cell function that can also contribute to alterations in neurologic function.

Neuroinflammation is also present in a variety of neurodegenerative diseases and is influenced by reactive microglia and astrocytes that lose their homeostatic functions. This leads to the release of proinflammatory cytokines, which can damage the blood-brain barrier (BBB) and lead to synaptic dysfunction, neurotoxicity, and neuronal death ^55^. In the context of COVID-19, CNS cells, including glial cells and neurons, have been identified as targets for SARS-COV-2 infection in both patients and animal models ^56–59^, but the virus is considered not to be highly neuroinvasive. Post-mortem analysis of brains from COVID-19 patients has reported gliosis, axonal damage, and BBB disruption^60–62^. Similar findings have been reported in laboratory animals, including mice and syrian hamsters. Infection with either SARS-CoV-2 or SARS-CoV-2-MA10 has led to BBB-disruption and gliosis across various anatomical regions including the hippocampus in the absence of direct viral infection of the CNS with decreased hippocampal neurogenesis, synaptic loss, as well as memory deficits ^63–66^. Cases where direct viral infiltration in the brain was observed, also noted similar results, including hippocampal gliosis. Additionally, an increase in expression of AD risk genes has been reported ^67^. Therefore, it is possible that COVID-19-associated neuroinflammation and specific neurodegenerative processes could result in hippocampal injury and neuronal/synaptic dysfunction as a result of infection.

In this study, we used infection of aged C57BL/6 (WT) and 5xFAD mice with a mouse-adapted SARS-CoV-2 [SARS-CoV-2 MA10 (MA10)] ^68,69^ to determine if infection is associated with neurologic changes in both strains of mice and to assess if there is an effect of infection on AD neuropathology. One of the most-widely used AD models are 5xFAD mice, which overexpress the Swedish, London, and Florida (I716V) mutations in human APP and M146L and L286V mutations in the PS1 gene ^70^. Extracellular plaques emerge by 2-months-old in the subiculum region of the hippocampus and as mice age, amyloid plaques progressively spread throughout the deep layers of the cortex and frontal cortex with corresponding neuroinflammation in the form of reactive astrocytes and activated microglia ^70,71^. Additionally, the presence of dystrophic neurites and impaired synaptic transmission trigger the onset of cognitive decline and neuronal loss by 9-months-old ^70,72^. Intranasal inoculation of both WT and 5xFAD mice, with MA10 resulted in similar acute lung pathology associated with increased expression of transcripts associated with inflammation and anti-viral immune responses, consistent with previous findings ^68^. We did not detect viral RNA in the CNS of either WT or 5xFAD mice yet bulk RNA sequencing and spatial transcriptomic analysis of the brains of both infected WT and 5xFAD mice indicated changes in pathways associated with neuronal dysfunction, astrocyte and microglial homeostasis, and reduced expression of adhesion molecules in vascular endothelial cells. MA10 infection of 5xFAD mice did not affect Aβ plaque volume or number, indicating that in this transgenic AD model, this pathological feature is not affected.

## Materials and Methods

### Viral infection of mice

Male and female 10–14-month-old C57BL/6 WT and 5xFAD were used for these studies. Tg (APPSwFlLon, PSEN1*M146L*L286V) 6799Vas/ Mmjax RRID:MMRRC034848-JAX, was obtained from the Mutant Mouse Resource and Research Center (MMRRC) at The Jackson Laboratory, an NIH-funded strain repository, and was donated to the MMRRC by Robert Vassar, Ph.D., Northwestern University. Mouse-adapted SARS-Related Coronavirus 2 MA10 Variant (in isolate USA-WA1/2020 backbone), Infectious Clone (ic2019-nCoV MA10) in Calu-3 Cells, NR-55329 developed by Baric and colleagues ^68^ was used for infection of experimental mice. Age-matched (10-14 months) 5xFAD and WT mice were intranasally inoculated with 5 x 10³ pfu of SARS-CoV-2 MA10 in 50µl of 1x PBS or sham inoculated. Infected and uninfected mice were monitored daily for signs of clinical illness.

### Histology and immunohistochemical staining

Mice were euthanized at defined times post-infection (p.i.) and brain and lungs were collected. Half brains were fixed in 4% PFA for 24 hours at 4 °C, then transferred into 30% sucrose in 1 X PBS solution for cryoprotection for 48 hours at 4 °C, embedded in O.C.T and 15µm sagittal sections were cut via cryostat (Thermo 95 664 OEC70 Micron HM525). Brain sections were desiccated overnight, and slides were rinsed with 1X TBS to remove residual O.C.T. Samples were blocked with 5% normal goat serum and 0.1% Triton-X in TBS, followed by overnight at 4 °C incubation with appropriate primary antibodies. On the second day, slides were treated with appropriate secondary antibodies following TBS rinsing. Amylo-Glo staining (TR-300-AG; Biosensis) was performed according to the manufacturer’s instructions. A complete list of antibodies and dilutions is provided in **Table 1**. Lungs were fixed and processed in the same manner as described above and stained with hematoxylin/eosin (H&E).

**Table 1:** Primary, secondary antibodies, and associated information.

| Primary antibody | Source | Catalog # | Dilution | Secondary antibody | Source | Catalog # | Dilution |
| --- | --- | --- | --- | --- | --- | --- | --- |
| IBA1 | Wako | 019-19741 | 1:500 | Goat anti-rabbit IgG, AlexaFluor 488 | ThermoFisher | A11008 | 1:200 |
| GFAP | Abcam | ab134436 | 1:1000 | Goat anti-chicken IgY Alexa Fluor 594 | Abcam | ab150172 | 1:400 |
| Mac2 | Cedarlane | CL8942AP | 1:500 | Goat anti-rat IgG Alexa Fluor 594 | Abcam | ab150160 | 1:200 |
| PSD95 | Abcam | ab18258 | 1:500 | Goat anti-rabbit IgG, AlexaFluor 488 | ThermoFisher | A11008 | 1:200 |
| Synaptophysin | Sigma-Aldrich | S5768 | 1:1000 | Goat anti-mouse IgG Alexa Fluor 555 | ThermoFisher | A21424 | 1:200 |
| NeuN | Abcam | ab104225 | 1:100 | Goat anti-rabbit IgG, AlexaFluor 488 | ThermoFisher | A11008 | 1:350 |
| LAMP1 | Abcam | ab25245 | 1:200 | Goat anti-rat IgG Alexa Fluor 594 | Abcam | ab150160 | 1:200 |

### Imaging and quantitative analysis

High-resolution fluorescent images were obtained at 20X magnification using a Leica TCS SPE-II confocal microscope and LAS-X software. With confocal imaging, one field of view (FOV) per brain region was captured per mouse. Three brain regions per mouse were imaged: cortex (CRTX), dentate gyrus (DG) and subiculum (SUB). Cell counts and plaque burden were quantified using the spots module in Imaris. Volumetric measurements were automatically acquired using the surfaces module with confocal images of each brain region. Super-Resolution Lattice Structure Illumination Microscopy (Lattice-SIM) was performed using an Elyra 7 microscope system. The same three distinct anatomical regions (CRTX, DG, and SUB) per mouse were imaged at 40X magnification. Images were processed using ZEN SIM2 on the ZEN software before being analyzed on Imaris. For quantification of both pre- and post-synaptic puncta (synaptophysin and PSD95), the Imaris spots module was used to count the number of puncta for each z-stack. Z-stacks were 5-8uM and the number of co-localized spots was normalized to z-stack volume. Overlapping puncta were defined as spots that were <0.2 uM apart.

### RNA extraction and PCR

Mouse tissue was added to TRIzol and homogenized with the Bead Ruptor 12 (Omni International) and 1.4mm ceramic beads (Omni-International, 19-627). RNA was extracted via RNeasy minikit (Qiagen, 74106) using the “Purification of Total RNA, Including Small RNAs, from Animal Tissues’’ protocol, with Buffer RWT substituted by Buffer RW1. Library preparation, RNA sequencing, and read mapping analysis were performed by Novagene Co. Gene expression values were normalized into log2 FPKM (fragments per kilobase of transcript per million mapped reads). Heatmaps were created using Morpheus (Morpheus, https://software.broadinstitute.org/morpheus). Volcano plots were created using custom code in R. Differentially expressed genes were selected as significant with a log2(Fold Change) > 0.5 and a false discovery rate (FDR) <0.05. From the comparisons, lists of genes of interest were chosen to plot a heatmap of their expression and a GO term enrichment analysis using enrichR (https://amp.pharm.mssm.edu/Enrichr/).

### cDNA synthesis and gene analysis

cDNA was made following the “First Strand cDNA Synthesis” protocol by New England Biolabs, using AMV Reverse Transcriptase (New England Biolabs, M0277L), Random Hexamers (Invitrogen, N8080127), RNAse inhibitor (New England BiolabsM0314L) and AMV Buffer (New England Biolabs B0277A). qPCRs were performed using the QIAquant 96 2 plex and iTaq Universal SYBER Green Supermix (BioRad, 1725120). Standard protocol for iTaq Universal SYBER GREEN Supermix was followed. Reactions were 10µl, and the machine was set to run for 1 cycle (95°C for 3 min), followed by 40 cycles (95°C for 15 s, then 58°C for 30 s). Sequences for mouse GAPDH, forward primer: AACTTTGGCATTGTGGAAGG, reverse primer: GGATGCAGGGATGATGTTCT; SARS-CoV-2 Nucleocapsid primers were purchased from Integrated DNA Technologies, forward (IDT, 10006821) and reverse (IDT, 10006822).

### Enzyme-linked immunosorbent assay

SARS-CoV-2 spike protein-specific antibodies in plasma of infected mice were measured by ELISA. 96-well plates were coated with 100 ng per well of Recombinant SARS-CoV (2019-nCoV) Spike S1+S2 ECD-His Recombinant Protein # 40589-V08B1 from Sinobiological and incubated overnight at 4°C. Plates were then washed three times with 1X PBS and blocked with 3% BSA in 1X PBS for 1 hour at room temperature. Following blocking, serial dilutions of mouse plasma (100 μl/well) were added and incubated for 2h at RT. Bound antibodies were detected using HRP-conjugated goat anti-Mouse IgG (H+L) Catalog # 31430 from Invitrogen and developed with TMB substrate (ThermoFisher, REF # 34028). The reaction then stopped with Stop reagent for TMB substrate catalog number S514-100ML from Millipore. Plates were read with the ELISA reader at 450nm.

### Cytokine profiling using Luminex platform

One aliquot of frozen plasma samples from mice were thawed to perform cytokine profiling using the Luminex Intelliflex platform. We first performed a titration assay, 3 samples from each cohort were used and diluted at 1:25, 1:50, and 1:100. Then, these three dilutions and a neat control were screened using the MILLIPLEX® MAP mouse kit “Milliplex Mouse Cytokine/Chemokine MAGNETIC BEAD Premixed 32 Plex Kit.”, which was the same kit intended for general assay. After measuring analytes and comparing concentrations, neat samples were chosen as the best representatives of concentration values in the samples measured. Hence, plasma samples were used in a neat state (no dilution) in the general assay and 10µl of each sample were assessed in a 384-well plate for cytokine profiling. The Milliplex Mouse Cytokine/Chemokine MAGNETIC BEAD Premixed 32 Plex Kit (G-CSF, EOTAXIN, GMCSF, IFNY, IL1a, Il1b, IL2, IL4, IL3, IL5, IL6, IL7, IL9, IL10, IL12P40, IL12P70, LIF, IL13, LIX, IL15, IL17, IP10, KC, MCP1, MIP1A, MIP1B, MCWF, MIP2, MIG, RANTES, VEGF, TNFA) was used to measure plasma cytokine and chemokines. To adjust the manufacturer’s protocol to our 384-well plate format, all other reagents, including antibodies, magnetic beads, and detection reagents, were used at 10 µL. Outcomes from wells with <50 beads for each analyte were excluded from analysis. Plates were read using a xMAP INTELLIFLEX® System (Luminex).

### Data analysis for Cytokine profiling

Quality control was performed on cytokine quantification data using the Belysa® Immunoassay Curve Fitting Software (Millipore). This consisted of evaluating standard curves for all 32 analytes and quality control samples (provided in each kit as QC1 and QC2) measured in the experiment. The interpolated concentrations of the plasma samples were exported in an excel file format for statistical analysis. Graph Pad Prism v. 10.5 was used for analysis and image generation. For statistical analysis a two-way ANOVA was fit to model the log expression of each cytokine. Significant differences were evaluated by group using the overall F test followed by pairwise comparisons under the Tukey’s Honest Significant Differences procedure.

### Spatial transcriptomic analysis

Formalin-fixed brain hemispheres were embedded in optimal cutting temperature (OCT) compound (Sakura Fintek, 4583) and stored frozen at −80°C. Sagittal sections were cut at 10um the day before starting the protocol on a Leica cryostat (LeicaBiosystems, CM1950). Six brain sections were mounted onto VWR Superfrost Plus slides (Avantor, 48311–703), allowed to dry at room temperature for 15 minutes to promote tissue adherence, and stored overnight at −80°C. Tissue was processed in accordance with the Nanostring CosMx fresh-frozen slide preparation manual for RNA (NanoString University). On day one of the slide preparation protocol, slides were removed from −80°C and baked at 60°C for one hour. Slides were then processed for CosMx: three 5-minute washes in 1X PBS, one 2-minute wash in 4% sodium dodecyl sulfate (Thermo Fisher Scientific, AM9822), : three 5-minute washes in 1X PBS, one 5-minute wash in 50% ethanol, one 5-minute wash in 70% ethanol, and two 5-minute washes in 100% ethanol before allowing slides to air dry at room temperature for 10 minutes. A pressure cooker was used to maintain slides in 100°C for 15 min in preheated 1X CosMx Target Retrieval Solution (Nanostring) for antigen retrieval. Slides were then cooled in DEPC-treated water (Thermo Fisher Scientific, AM9922) for fifteen seconds, transferred to 100% ethanol for three minutes, and then isolated at room temperature for 30 minutes to air dry. For effective tissue permeabilization, slides were transferred to a digestion buffer (3 μg/mL Proteinase K in 1X PBS; Nanostring) to incubate, then washed twice for five minutes in 1X PBS. Fiducials for imaging were diluted to 0.00015% in 2X SSC-T and left for five minutes on slides to incubate. Slides were shielded from light following fiducial application. A 10% neutral buffered formalin solution (NBF; CAT#15740) was applied for one minute to post-fix tissues. Slides were then washed for five minutes with NBF Stop Buffer (0.1M Tris-Glycine Buffer, CAT#15740) twice and washed for five minutes with 1x PBS. An NHS-Acetate (100 mM; CAT#26777) solution was then administered to the slides and incubated for fifteen minutes at room temperature. Slides were then washed twice for five minutes with 2X SSC and incubated in modified 1000-plex Mouse Neuroscience panel (Nanostring) with the addition of an rRNA segmentation marker for in situ hybridization in a 37°C hybridization oven overnight for 16–18 hours. On day 2 of slide preparation, slides were placed in preheated stringent wash solution (50% deionized formamide [CAT#AM9342]) in 2X saline-sodium citrate [SSC; CAT#AM9763]) at 37°C twice for 25 minutes each. Slides were then washed in 2X SSC twice for two minutes each. Slides were then incubated with DAPI nuclear stain for fifteen minutes, washed for five minutes with 1X PBS, incubated with histone and GFAP cell segmentation markers for one hour, and washed 3 times for five minutes each in 1X PBS. Adhesive flow cells (Nanostring) were placed in each slide to create a fluidic chamber for spatial imaging. Slides were loaded into the CosMx instrument and automatically processed. On each slide, approximately 600 total fields of view (FOVs; ∼100 FOVs per brain section) were chosen that captured hippocampal, corpus callosum, upper thalamic, upper caudate, and cortical regions for each of the six sections. Imaging progressed for approximately seven days and data were uploaded automatically to the Nanostring AtoMx online platform. Pre-processed data was exported from AtoMx as a Seurat object for analysis. Spatial transcriptomics datasets were processed with R 4.3.1 software as previously described. Principal component analysis (PCA) and uniform manifold approximation and projection (UMAP) analysis were performed to reduce dataset dimensionality and visualize clustering. Data-driven clustering at 1.0 resolution yielded 46 clusters. Clusters were manually annotated based on transcript expression of known marker genes and location in XY space. Cell proportion plots were generated by first visualizing the number of cells in each cell type, then scaling relative to 1) normalized percentages per group, calculated by dividing the number of cells in each cell type-group pair by the total number of cells, and 2) dividing by the sum of the proportions across the cell type to account for differences in sample sizes. MAST was used on scaled expression data to perform differential gene expression analysis per cell type between groups to compute the average difference, defined as the difference in log-scaled average expression between the 2 groups for each non-specific cell type. The absolute log_2_ fold change values of all genes with statistically significant gene (i.e., padj < 0.05) differential expression patterns between two groups were summed to calculate the DEG scores between group pairs for each subcluster. ggplot2 3.4.4174 was used to generate data visualizations.

### Statistical analysis

GraphPad Prism was used to perform statistical analysis on immunohistochemical data. Data were analyzed using two-way ANOVA, and Tukey’s post-hoc tests used to evaluate biologically relevant interactions. Each data set was examined for sex-differences, and if none were found, both male and female data were combined; p-value of <0.05 was considered statistically significant. Data for each experiment is presented as mean±SEM.

## Results

### MA10 infection of WT and 5xFAD mice

C57BL/6 (WT) and 5xFAD mice (10-14 months of age) were intranasally (i.n.) infected with 5×10³ pfu of MA10. Animals were sacrificed at days 7 and 21 (p.i.) and lungs and brains harvested to assess tissue pathology, presence of viral RNA, and effects on AD-associated neuropathology. By 4 months of age, 5xFAD mice develop Aβ plaques in the brain which increase in number and size over time, plateauing between 10-12 months of age ^70,73^. By day 21 p.i., there was no significant difference in mortality, with ∼20% of infected 5xFAD mice and ∼15% of infected WT mice dying in response to infection (**Figure 1A**). Daily weighing of mice showed MA10 infected WT mice exhibited significant weight loss compared to their uninfected controls only at 1 and 5-7 p.i., while MA10 infected 5xFAD mice did not demonstrate significant weight loss in comparison to their uninfected controls (**Figure 1B**). Spike-specific IgG antibodies were detected at days 7 and 21 p.i. in both WT and 5xFAD mice confirming infection with MA10 compared to SHAM-infected mice (**Figure 1C**). Viral RNA was examined via qPCR in lungs and brains of infected mice at 7 days p.i. Low levels of viral RNA were present at day 7 p.i. within the lungs (**Figure 1D**). No viral RNA was detected in the brains at 7 days p.i. (**Figure 1E**) consistent with previous reports indicating MA10 does not readily infect the CNS ^63^. There were no significant differences between uninfected and infected cohorts, and any Ct values yielded from infected cohorts in qPCR were assumed to be background, as they were like those of uninfected/control mice. Hematoxylin and eosin (H&E) staining of lungs of infected WT and 5xFAD mice at day 7 p.i. demonstrated both alveolar and interstitial lesions, with alveolar hemorrhage and edema, interstitial congestion, and lymphocytic infiltrates within the lungs and by day 21 p.i. there was still evidence of inflammation in both MA10-infected WT and 5xFAD mice although this was reduced in comparison to day 7 p.i. (**Figure 1F**). Examination of proinflammatory cytokines and chemokines within the blood of MA10-infected WT and 5xFAD mice at day 7 p.i. showed a trend toward increased levels of proinflammatory cytokines in infected mice compared to their controls, however, the differences did not reach statistical significance for all markers. MA10 infection did result in a significant increase in CCL5 levels in both WT (p<0.001) and 5xFAD (p<0.05) mice compared to uninfected controls while only CCL2 was significantly (p<0.01) increased in infected 5xFAD compared to uninfected 5xFAD mice (**Figure 1G**).

**Figure 1.**
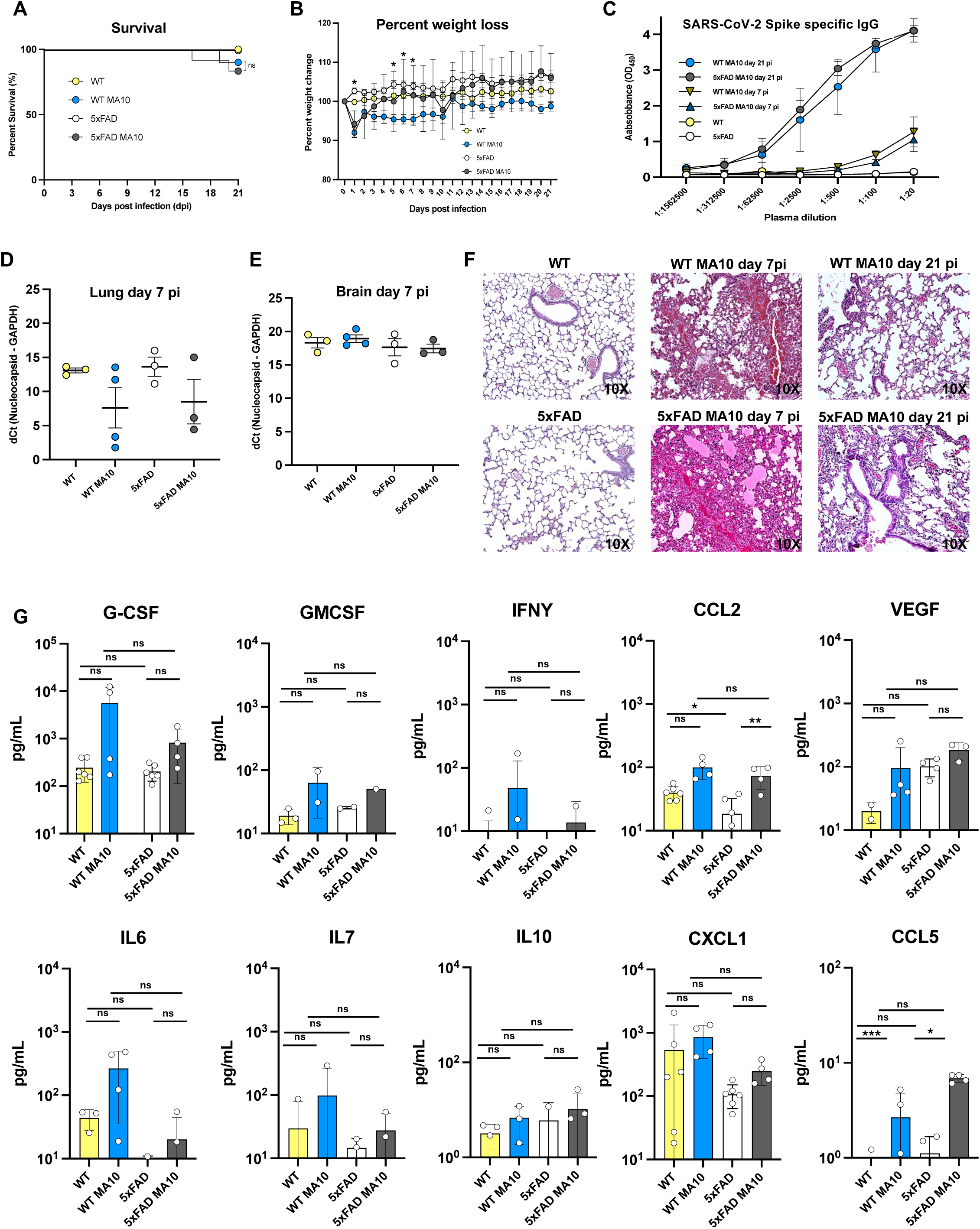
MA10 infection elicits mild disease in infected mice, and genotype does not affect mortality. (**A**) Percent survival of uninfected WT and 5xFAD compared to MA10 infected WT and C57BL/6 mice. (**B**) Average body weight changes each day p.i normalized to the body weight on the day of infection. (**C**) Neutralizing IgG anti-spike antibodies were present in the plasma of MA10-infected mice at days 7 and 21 p.i. as assessed by ELISA. RT–qPCR was performed on (**D**) lung and (**E**) brain homogenates at day 7 p.i. to quantify MA10 nucleocapsid (N) gene expression. The data are presented as dCt values, calculated by subtracting the Ct of the housekeeping gene GAPDH from the Ct of the viral N gene. Lower dCt values indicate higher viral RNA abundance. Each dot represents an individual mouse; bars represent the mean ± SEM. (**F**) Representative H&E-stained lung images at 10× magnification from uninfected and MA10-infected WT and 5xFAD mice at both 7 and 21 days p.i., are shown. Pathological features associated with infection included airway edema, vascular congestion, and intra-alveolar hemorrhage, peri-bronchiolar lymphocytic cuffing, and interstitial vascular congestion and lymphocytic infiltrates. (**G**) Cytokine concentrations in plasma were measured using a multiplex bead-based immunoassay on the Luminex platform. Data for individual cytokines are shown with each dot representing an individual mouse; in some cases, specific cytokines in mice were below level of detection. Data analyzed using two-way ANOVA and presented as mean ± SEM; *p<0.05; **p<00.01.

### Microglia and astrocytes in MA10-infected mice do not exhibit changes in numbers nor volume

Brains from MA10-infected and control mice were immunofluorescently labeled for microglia (IBA1) and astrocytes (GFAP) to determine if there was an increase in glial activation in response to MA10 infection at day 21 p.i. We elected to examine the cortex (CRTX), dentate gyrus (DG), and subiculum (SUB) as previous studies of 5xFAD mice indicate changes in AD associated pathology in these regions ^73^. Additionally, laboratory animals with SARS-CoV-2 or MA10 indicated CNS changes within these anatomical regions as a result of infection ^63–67^. As a metric of cell activation, we measured either GFAP+ or IBA1 cell numbers and volume in control and MA10-infected WT and 5xFAD mice. Representative images of IBA1+ microglia and Mac2+ monocytes along with Amyloglo counterstain for Aβ plaques (5xFAD mice) were acquired within the CRTX, DG, and SUB from uninfected and MA10-infected WT and 5xFAD mice are shown (**Figure 2A**). Quantification of microglia numbers (**Figure 2B**), and volume (**Figure 2C**) does not indicate significant changes in MA10-infected WT or 5xFAD mice compared to their uninfected controls. There was an increase in microglia number and volume in both uninfected and infected 5xFAD mice compared to uninfected and infected WT mice in all brain regions examined (**Figure 2C**). MAC2 is a unique marker for peripheral monocyte-derived macrophages that allows discrimination between these cells and microglia ^74^. Several studies have shown that under inflammatory conditions, peripherally derived monocytes enter the brain in response to infection ^74–76^. Immunostaining and quantification do not indicate changes in MAC2-positive cell number or volume within the cortex, dentate gyrus, or subiculum as a result of MA10 infection in either WT or 5xFAD mice (**Supplemental Figure 1**). There were increased numbers (p<0.0001) of MAC2-positive cells in the brains of uninfected 5xFAD mice compared to uninfected WT mice, reflective of the neuroinflammation occurring within the brains of the 5xFAD mice as a result of ongoing AD neuropathology and Aβ plaque accumulation (**Supplemental Figure 1**). Immunofluorescent staining of GFAP+ astrocytes in uninfected and MA10-infected WT and 5xFAD mice along with Amylglo staining of Aβ plaques in control and MA10-infected 5xFAD mice within the CTX revealed no differences in GFAP expression (**Figure 2D**). Quantification of both astrocyte numbers and volume revealed no differences between control and infected WT and 5xFAD mice, indicating that infection did not result in increased astrocyte activation at 21 days p.i. (**Figures 2E-F**). There were increased numbers of GFAP+ cells within the cortex of uninfected 5xFAD mice compared to uninfected WT mice while volume of GFAP+ cells was increased in both the cortex and subiculum in uninfected 5xFAD mice compared to uninfected WT mice (**Figure 2F**). GFAP numbers were reduced in the dentate gyrus of uninfected 5xFAD mice compared to uninfected WT mice (**Figure 2F**). Overall, these findings indicate that MA10 infection of either WT or 5xFAD does not result in sustained activation of either microglia or astrocytes by 21 days p.i.

**Figure 2.**
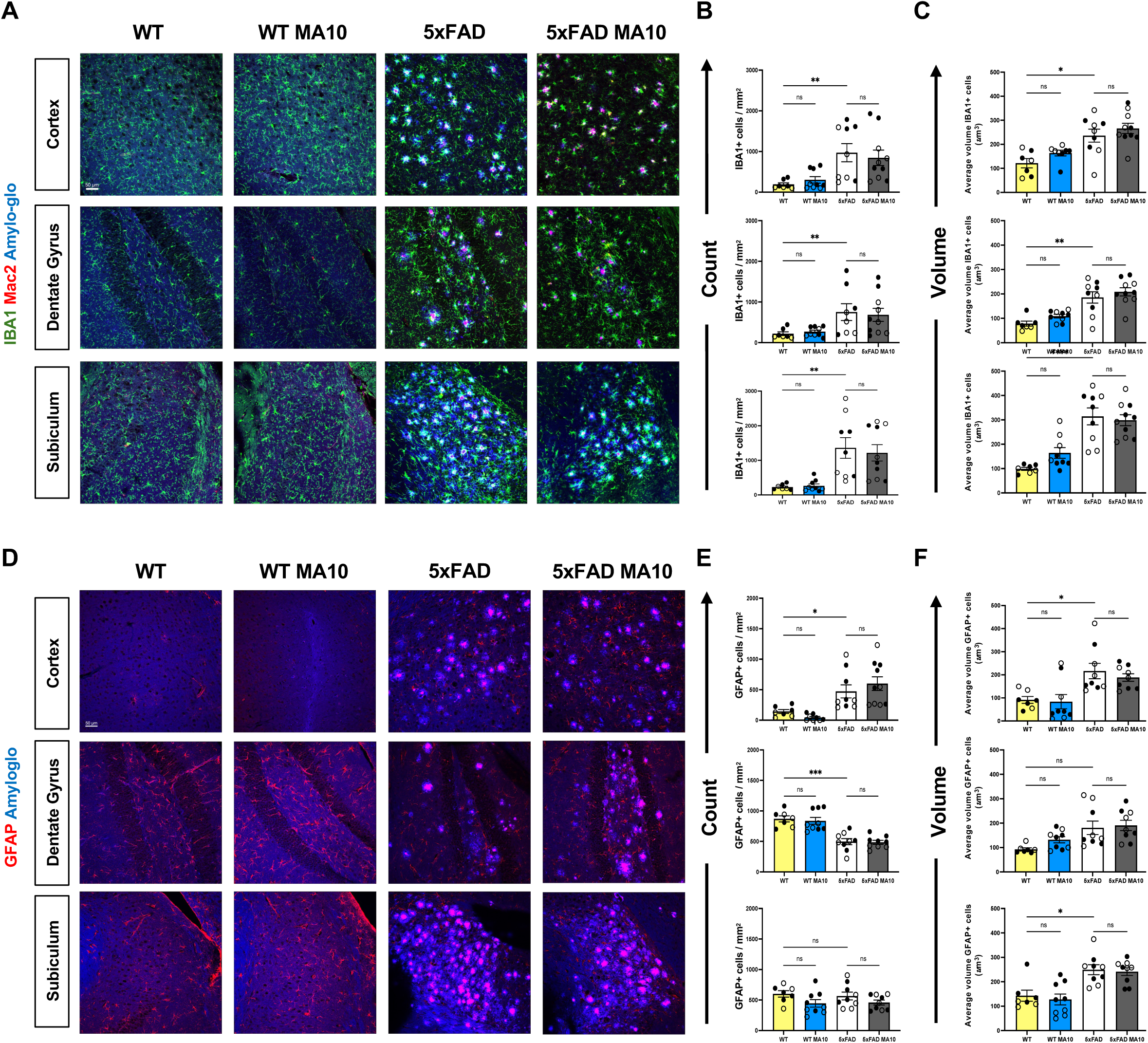
Effect of MA10 infection on glial activation in WT and 5xFAD mouse brains. Brains from uninfected and MA10-infected WT and 5xFAD mice at day 21 p.i. were immunostained for IBA1 and MAC2 to assess changes in microglia and infiltrating monocytes/macrophages, respectively and GFAP to detect astrocytes. Dense-core Aβ plaques were stained via Amylo-Glo. (**A**) Representative images (20X magnification) from brains isolated from uninfected and infected WT and 5xFAD mice showing the cortex, dentate gyrus, and subiculum stained for microglia, macrophages, and Amylo-Glo. (**B**) Average IBA1+ cells per mm^2^ in cortex (upper graph), dentate gyrus (middle graph) and subiculum (bottom graph) from experimental mice. (**C**) IBA1 average volume per um^3^ in cortex (upper graph), dentate gyrus (middle graph) and subiculum (bottom graph) from experimental mice. (**D**) Representative images from brains isolated from uninfected and infected WT and 5xFAD showing GFAP expression and Aβ plaques. Average GFAP+ cells per mm^2^ (**E**) and average volume per um^3^ (**F**) in the cortex (upper graphs), dentate gyrus (middle graphs) and subiculum (bottom graphs). Immunohistological data were analyzed using two-way ANOVA. Tukey’s post-hoc test was employed to examine biologically relevant interactions. Female and male mice are indicated by open or closed circles respectively. Data are represented as mean ±SEM; * p < .05, ** p < .01, *** p < .001. Scale bar in (A) = 50 µm.

### Dystrophic neurite and Aβ plaque burden in MA10-infected mice

Dense core Aβ plaques, a key feature of AD pathology, are surrounded by dystrophic neurites, which are abnormal swollen neuritic processes that can be detected by immunostaining for lysosome-associated membrane protein 1 (LAMP1)^73^. Representative images of LAMP1+ lysosomes with Amylo-glo-positive Aβ plaques were acquired for the cortex, dentate gyrus, and subiculum (**Figure 3A**). Dystrophic neurites and Aβ plaque pathology are only present in 5xFAD, and infection does not affect or induce these pathologic features in WT mice (**Figure 3A**). Immunostaining and quantification do not indicate changes in count or volume of dystrophic neurites in response to MA10 infection of either WT or 5xFAD mice compared to uninfected controls in any of the brain regions examined in either WT or 5xFAD mice (**Figures 3B-C**). There were no differences in either Aβ plaques count or volume in MA10 infected 5xFAD mice compared to uninfected 5xFAD mice (**Figure 3D-E**). Synaptic puncta were next quantified via co-localization of the presynaptic marker, synaptophysin, and postsynaptic marker PSD95 to assess effects of MA10 infection on neuronal dysfunction. Representative images of synaptophysin puncta, and PSD95 positive puncta were acquired (**Supplemental Figure 2A**). Immunostaining and quantification did not indicate changes in synapse numbers in the brains of either WT or 5xFAD mice at 21 dpi (**Supplemental Figure 2B**).

**Figure 3.**
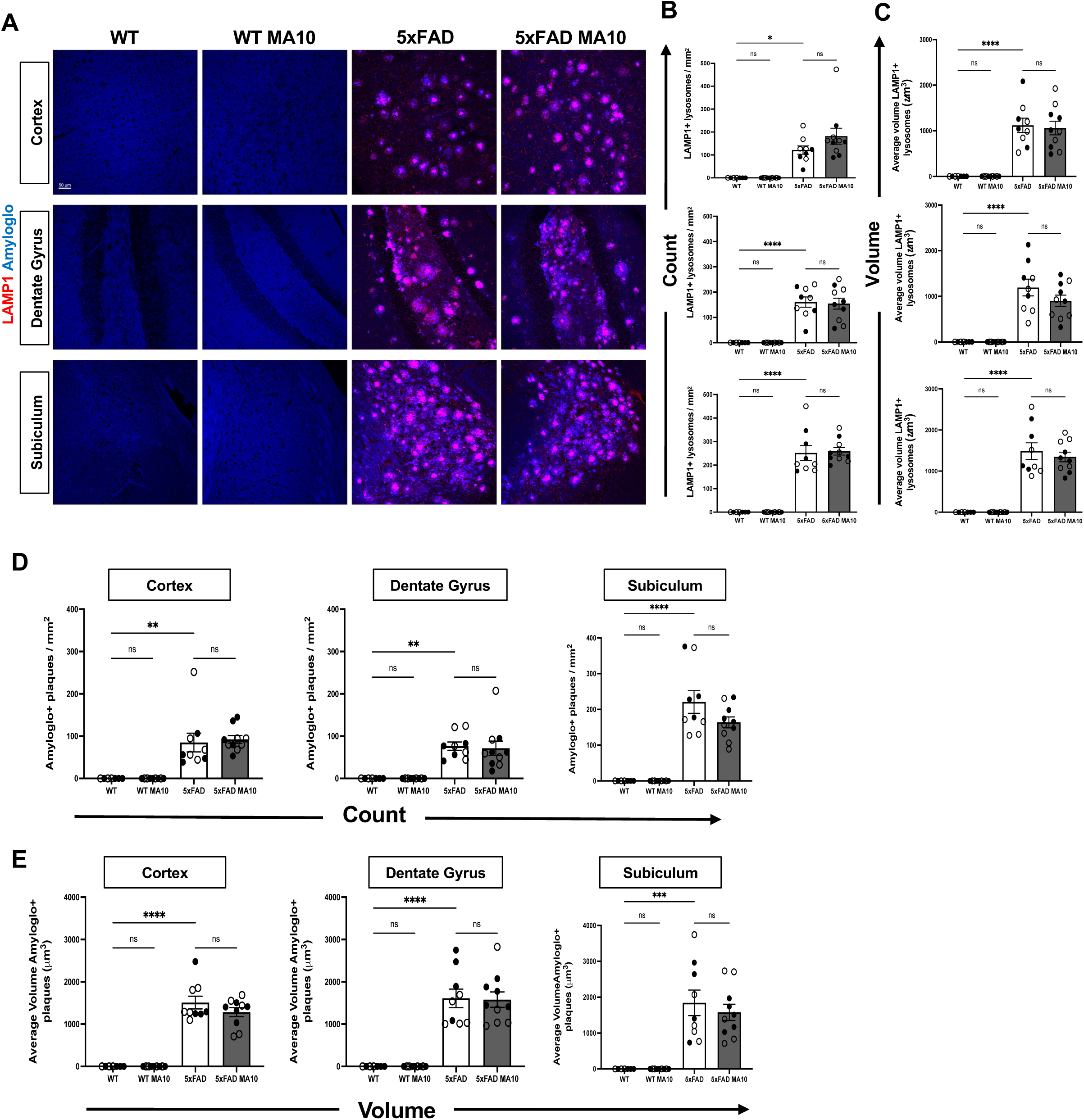
MA10 infection does not affect dystrophic neurites or Aβ plaques in 5xFAD mice. Brains from uninfected and MA10-infected mice at 21 days post-infection were immunostained with LAMP1 to visualize dystrophic neurites and with Amylo-Glo to detect dense-core Aβ plaques. (**A**) Representative images of LAMP1 and Amylo-Glo staining in the cortex, dentate gyrus, and subiculum of uninfected and MA10-infected WT and 5xFAD mice. **(B)** Quantification of LAMP+ lysosomes density (**B**) and volume (**C**) shows no differences in MA10-infected WT or 5xFAD mice compared to uninfected controls. Aβ plaque burden was determined through quantifying Amylo-Glo⁺ plaque number as shown in (**D**) and summarizes the average volume of these plaques (**E**). Immunohistological quantifications were analyzed using two-way ANOVA followed by Tukey’s post hoc test. Female and male mice are represented by open and closed circles, respectively. Data are presented as mean ± SEM; *p≤0.05, **p≤0.01, ***p<0.001, ****p≤0.0001. Scale bar in (A) = 50 μm.

### Bulk RNA sequencing of brains from MA10-infected mice

Bulk RNA sequencing of differentially expressed genes was performed on the brains of experimental mice at day 7 and 21 p.i. Volcano plots of brain gene expression in MA10 infected and control WT and 5xFAD at days 7 (**Figures 4A**) and 21 p.i. (**Figures 4E**) were plotted. Differentially expressed genes were selected as significant with a log2(Fold Change) > 0.5 and a false discovery rate (FDR) <0.05. On both days 7 and 21p.i., infected WT and 5xFAD groups show differential expression of genes when compared to control groups. At day 7 p.i, both groups exhibit a strong transcriptional response, with numerous genes significantly upregulated (**Figure 4A**). By day 21p.i, MA10-infected WT mice showed fewer numbers of DEGs compared to MA10-infected 5xFAD mice, suggesting a more persistent molecular response in MA10-infected 5xFAD mice (**Figure 4E**). Heatmaps were created to compare significant DEGs from control and MA10-infected WT and 5xFAD mice. DEGs were selected as significant if Log₂Fold Change was greater than 0.5 and false discovery rate/ adjusted p-value as less than 0.05 (FC>0.5, FDR<0.05). Genes that met such criteria were then used to plot heatmaps of their expression from the brains of experimental mice at days 7 p.i. (**Figure 4B**) and 21p.i. (**Figure 4F**). GO analysis of MA10-infected WT at day 7 p.i. revealed increased expression of pathways associated with voltage-gated calcium channel activity, abnormal operant conditioning, reduced long-term potentiation, and abnormal hippocampal mossy fiber morphology (**Figure 4C**). In the MA10-infected 5XFAD at day 7 p.i. analysis indicated changes in pathways associated with abnormal postsynaptic currents, synaptic transmission, neuron morphology, and spatial learning as well as impaired coordination and increased anxiety-related responses. Additionally, there are changes in pathways associated with synaptic transmission and plasticity as well as axonogenesis (**Figure 4D**). Brains of both the WT and 5xFAD groups did indicate lasting changes in gene expression (**Figure 4E**); however, heatmap analysis indicated that the MA10 infected 5xFAD group had dramatically more lasting differentially expressed genes in comparison to the MA10 infected WT group (**Figure 4F**). Moreover, when examining pathways of interest, MA10-infected WT mice showed only one identified down-regulated gene (*UQCRFS1*) that is associated with AD^77^. (**Figure 4G**). Analysis on the 5xFAD group at 21 days p.i. indicated changes of many more pathways, most of which were associated with neuronal and synaptic dysfunction, impaired coordination, and anxiety-related responses (**Figure 4H**). Furthermore, the brains of MA10-infected WT mice indicated changes in gene expression in pathways associated with abnormal sphingomyelin levels and axon guidance receptor activity (**Figure 4G**) while the brains of MA10-infected 5xFAD mice indicated changes in pathways associated with edema and abnormal circadian cycles (**Figure 4H**).

**Figure 4.**
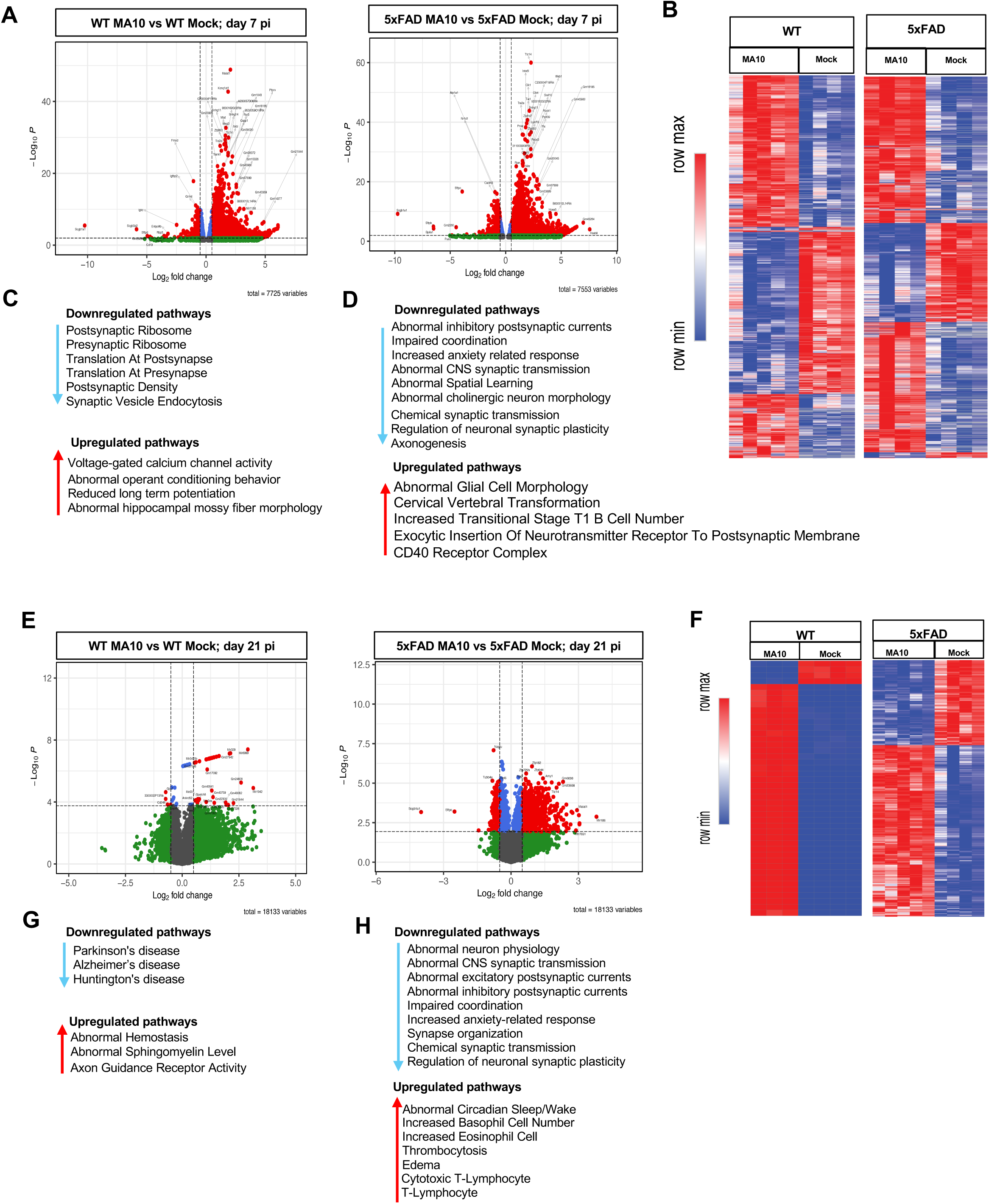
SARS2 MA10 infection in brains of WT and 5xFAD mice is associated with differential expression of genes involved in regulation of synaptic activity at both days 7 and 21 p.i. RNA sequencing was performed on brains from MA10 infected and control WT and 5xFAD male mice at 7 days p.i. (**A**) Volcano plot of differentially expressed genes (DEG) displaying fold change of genes (log2 scale) and P values. (**B**) Heatmap of selected (FC=0.5, FDR<0.05) DEGs from MA10-infected WT and 5xFAD and uninfected (mock) controls at day 7 p.i. Enriched pathways of interest examined via Gene Ontology (GO) expressed in (**C**) MA10-infected WT and (**D**) 5xFAD mice showing downregulated and upregulated pathways at day 7 p.i. Volcano plot of differentially expressed genes (DEG) displaying fold change of genes (log2 scale) and P values in (**E**) MA10-infected WT and 5xFAD mice at day 21 p.i. (**F**) Heatmap of selected (FC=0.5, FDR<0.05) DEGs from MA10-infected WT and 5xFAD and uninfected (mock) controls at day 21 p.i. Enriched pathways of interest examined via Gene Ontology (GO) expressed in (**G**) MA10-infected WT and (**H**) 5xFAD mice showing downregulated and upregulated pathways at day 21 p.i.

### Altered gene expression in resident CNS cells following MA10 infection of WT and 5xFAD mice

WT and 5xFAD MA10-infected and uninfected control brains were sectioned and processed for spatial transcriptomic analysis and analyzed for 1,000 genes using the Nanostring CosMx Spatial Molecular Imager platform (n = 3 per group, 1216 total FOVs, ∼100 FOVs per brain section) (**Figure 5A**; examples of cell segmentation in **Supplemental Figure 3**). 787,580 cells were captured with a mean transcript count of ∼600 per cell. Unbiased cell clustering identified 46 transcriptionally distinct cell clusters (**Figure 5B and Supplemental Figures 4A and B**), which were manually annotated based on gene expression and location. Projection of clusters in XY space revealed distinct localization to specific anatomical regions (**Figure 5C and Supplemental Figure 5)**. Separation of cell clusters by group (WT control, WT MA10 infected, 5xFAD control, 5xFAD MA10 infected) revealed distribution of distinct cell populations within the brains of each experimental group (**Figure 5D**). Clusters from the same broad cellular subtype were then combined into broad cell types (i.e. Astrocyte 1, Astrocyte 2, etc. were assigned to the Astrocyte cell type) for subsequent analyses. The proportion of cells in each major cluster from experimental groups was then plotted per genotype (**Figure 5E**). To broadly assess overall deviation from WT in a given group, DEG scores were calculated for each cellular subtype. To calculate DEG scores for each group, the log_2_fold change values were summed for DEGs that were significantly upregulated or downregulated in comparison to WT (i.e., p_adj_ ≤ 0.05), yielding DEG scores for each subtype. Subtypes were then plotted in XY space colored by DEG score for upregulated and downregulated genes to give a visual impression of the spatial distribution of DEGs and overall degree of deviation from WT control in the three experimental groups (**Figure 5F**). As expected, the 5xFAD MA10 infected brains had the highest upregulated and downregulated DEG scores, indicative of broad deviation from WT gene expression patterns. As disease-associated microglia (DAMs) are considered important in Aβ pathology in 5xFAD-infected mice ^78^, we next plotting the DAM cluster in XY space revealing that DAMs are distributed in both uninfected and MA10-infected 5xFAD mice in plaque-laden areas i.e., the subiculum of the hippocampus, throughout the cortex (**Figure 5G**).

**Figure 5.**
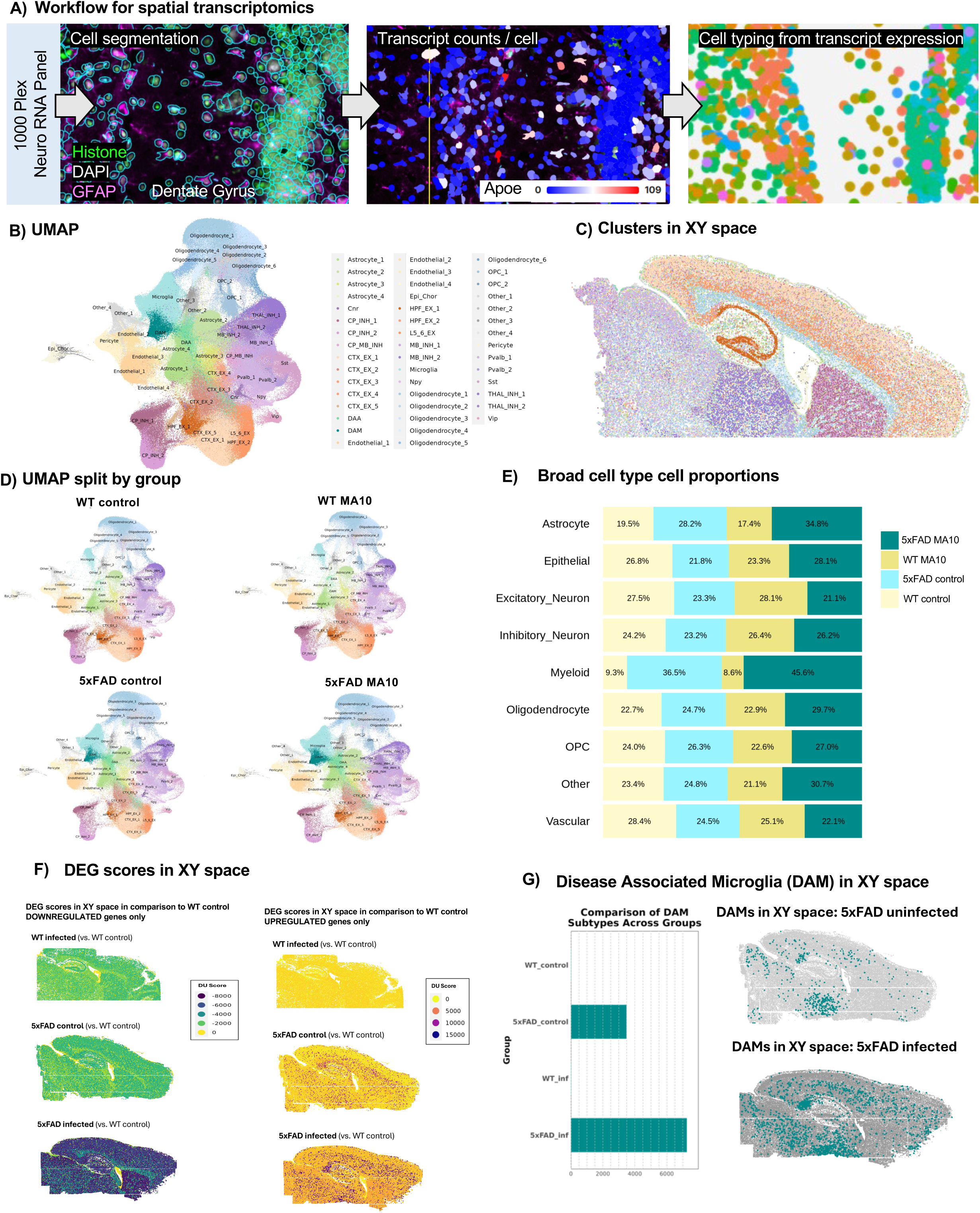
Spatial transcriptomic analysis of the MA10-infected WT and 5XFAD mouse brain. **(A)** Experimental workflow for targeted 1000-plex single-cell spatial transcriptomics. Fields of view (FOVs) were selected in hippocampal, corpus callosum, upper thalamic, upper caudate, and cortical regions for each of the six sections then imaged with DNA, rRNA, Histone, and GFAP markers for cell segmentation. Transcript counts for each gene were acquired per cell. (**B**) Uniform Manifold Approximation and Projection (UMAP) of 787,580 cells across 6 brains were captured with a mean transcript count of ∼600 per cell. Unbiased cell clustering identified 46 transcriptionally distinct cell clusters which were annotated with a combination of automated and manual approaches with reference to Allen Brain Atlas single-cell RNA-seq cell types, gene expression, and anatomical location in space. (**C**) 46 clusters plotted in XY space. (**D**) Separation of clusters by group (WT control, MA10-infected WT, 5xFAD control, MA10-infected 5xFAD). (**E**) Proportional distributions across experimental groups. Bar plots display the relative abundance of each cluster in each group. (**F**) Downregulated DEG scores in XY space in comparison to WT control (left panel), upregulated DEGs (right panel). (**G**) DAM cell counts across groups (left panel), distribution of DAMs plotted in XY space in uninfected and MA10 infected 5xFAD.

### MA10 infection alters gene expression in glial cells and neurons

Differential gene expression analysis revealed that MA10 infection led to the appearance of differentially expressed genes (DEGs) within broad cell types in both WT and 5xFAD mice, including microglia, astrocytes, oligodendrocytes, inhibitory neurons and vascular endothelial cells (**Figures 6A-J**). Many DEGs were identified between infected and uninfected control brains with infection in both 5xFAD and WT mice. In both mice, microglia demonstrated loss of numerous homeostatic signature genes^79^ (i.e. *Hexb, Csf1r, P2ry12, Cx3cr1*) (**Figures 6A and B**). Astrocytes indicated a downregulation of *Slc1a2, Gja1, Plpp3, Atp1a2, Ptprz1*, genes identified to be involved in astrocytic functions that support neuronal communication and homeostasis ^80–82^ (**Figures 6C and D**). Furthermore, oligodendrocytes showed decreased expression of genes related to myelination, lipid metabolism, and neuroprotection *(Malat1, Mag, Ugt8a,* and *Plp)*^83–85^ as well as an increased expression in genes related to cellular stress responses (*Sgk1)* ^86^(**Figures 6E and F**). Other cell types such as inhibitory neurons exhibited a downregulation of genes associated with neurotropic support (*Ndrg4, Sst, and Npy*)^87–92^ (**Figures 6G and H**), while vascular endothelial cells indicated consistent changes across genotypes in both the downregulation of genes associated with blood brain barrier (BBB) integrity (*Bsg, Cldn5, and Pecam1*)^93,94^ as well as the upregulation of genes associated with other vascular adaptations (*Apod, and Acta2*)^95,96^ (**Figures 6I and J**).

**Figure 6.**
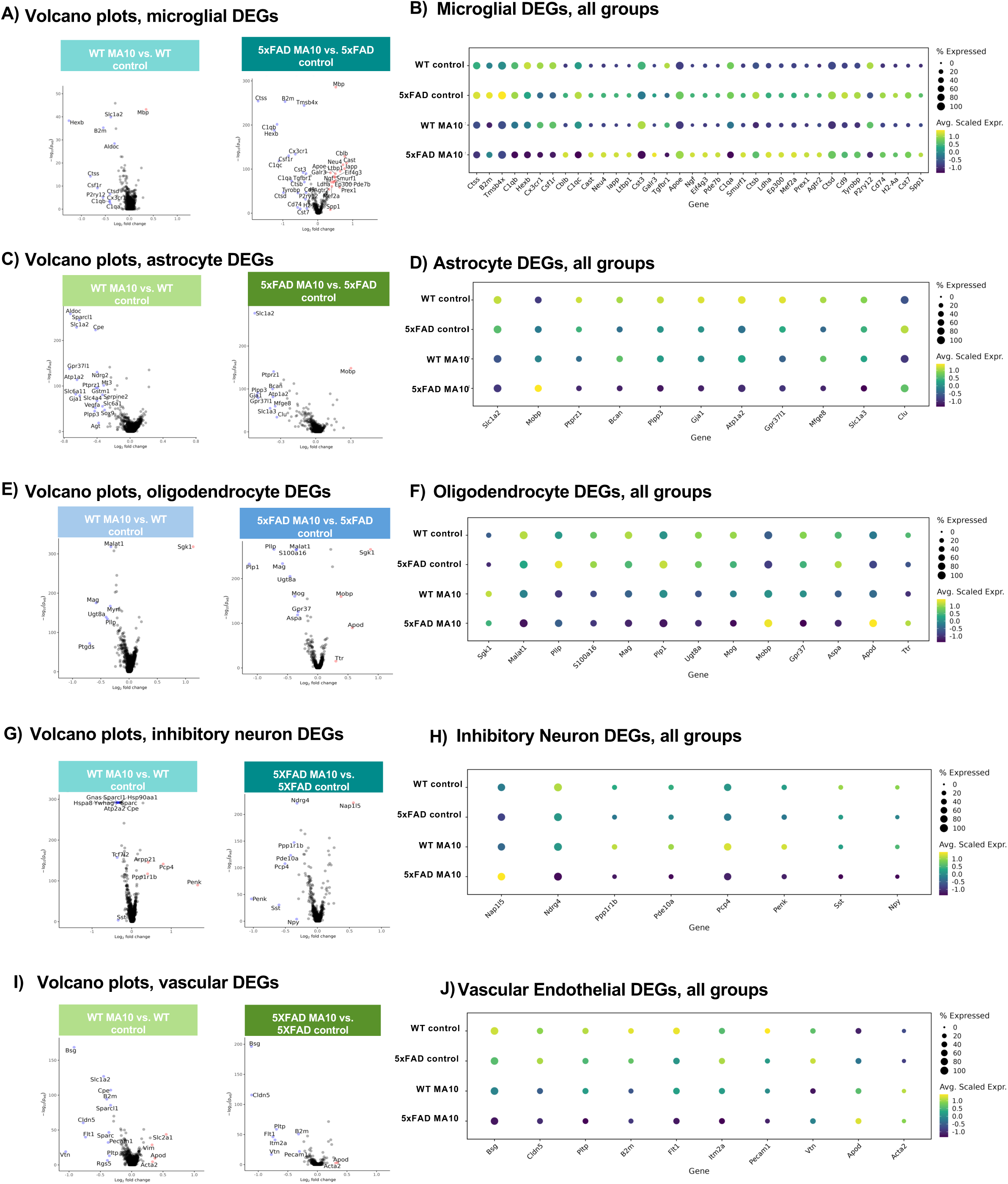
SARS2 MA10 infection leads to changes in gene expression within resident CNS cells, as revealed by spatial transcriptomic analysis of WT and 5xFAD mouse brains. Volcano plots of DEGs between each of the following broad cell type: (**A**) microglia, (**C**) astrocyte (**E**) oligodendrocyte, (**G**) inhibitory neuron and (**I**) vascular of both MA10-infected WT vs WT control (left panel) and MA10-infected 5xFAD MA10 vs 5xFAD control (right panel). (**B**), (**D**), (**F**), (**H**) and (**J**) represent corresponding dot plots showing the expression of individual genes across experimental groups (WT control, 5xFAD control, WT MA10, and 5xFAD MA10). Dot size represents the percentage of cells expressing the gene, while color indicates the average scaled expression level.

WT brains demonstrated a greater number of significant DEGs and/or a greater average difference than 5xFAD brains between infected and uninfected controls in the majority of broad cell types. Microglia represented a notable exception in that they were far more affected in 5xFAD brains than WT brains following infection; this most likely is that 5xFAD microglia are already highly dysregulated as a result of Aβ accumulation over time. While both WT and 5xFAD genotypes showed loss of homeostatic microglial markers after infection in comparison to their respective uninfected controls, the average difference (log_2_fold change) and significance (p_adj_) in 5xFAD microglia were higher than WT. In addition to conservation of the majority of downregulated genes from WTs, 5xFAD microglia also had a number of upregulated genes that were not present in WTs, including *Cblb, Cast, Neu4, Iapp, Apoe, Ltbp1, Eif4g3, Ngf,* and *Smurf1*, among others (**Figure 6B),** indicating a further loss of microglial homeostatic functions such as survival, communication, migration and phagocytic capacity to an even more reactive state suggesting that 5xFAD microglia are more affected by MA10 infection than WT microglia at the transcriptomic level.

### Comparing effects of Aβ pathology and MA10 infection on cell-type specific differential gene expression

We have demonstrated that MA10 infection in the periphery shifts the transcriptional state of microglia towards a greater loss of homeostatic markers. To investigate whether these dysregulated responses in microglia are a consequence of peripheral MA10 infection, the presence of Aβ pathology, or unique to the presence of both inflammatory stimuli, we compared the average differences in gene expression within all microglia in our spatial transcriptomic dataset. We plotted all DEGs found in comparing control 5xFAD vs. control WT samples (i.e., response to Aβ-associated pathology) and those found comparing MA10 infected WT vs. control WT (i.e., response to MA10 infection) using the average difference of each gene among the two comparisons (**Figure 7A**). In response to either Aβ pathology or MA10 infection, microglia shared only four significantly down-regulated DEGs (*Csf1r, P2ry12, Cx3cr1, Hexb)*, all of which are signature microglial homeostatic markers (**Figure 7B**). Meanwhile, there were several genes that were uniquely differentially expressed in one inflammatory stimulus over the other. Specific to MA10 infection, microglia demonstrated seven significantly down-regulated genes *(Ctsd, B2m, Ctss, C1qb, Aldoc, Slc1a2, and C1qa).* In contrast, microglia demonstrated 60 significantly up-regulated genes (*Apoe, Cd74, Cst7, Ctsd, Ctsb, Cd9, Clec7a etc.*) and 12 significantly down-regulated genes (*Selplg, Serinc3, Tmem119, Hmgb1, Camk2a, Malat1, Tgfbr1, etc.)* that were unique to responses to Aβ pathology (**Figure 7B**).Therefore, while the microglia activation response to peripheral MA10 infection and Aβ pathology down-regulated similar homeostatic markers, this comparison highlights the nuanced differences in related groups of genes when considering the magnitude of their inflammatory response.

**Figure 7.**
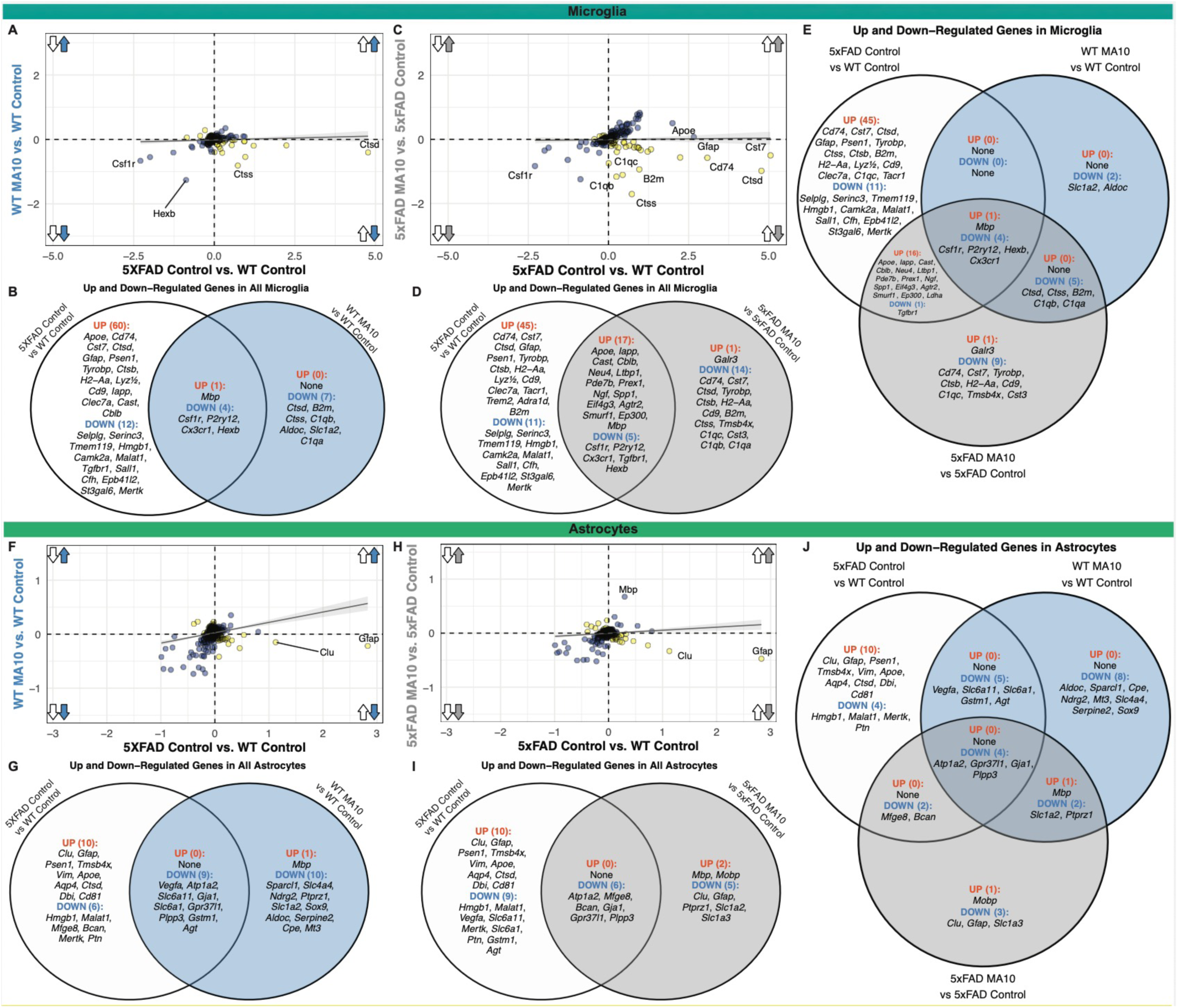
Microglia and astrocytes have differential responses to the presence of amyloid pathology, peripheral MA10 infection, or both concurrently. (**A**) Scatterplot of the average difference of all microglia for all significant genes (padj < 0.05) between 5xFAD control vs. WT control (x-axis, only amyloid pathology) and WT MA10 vs. control WT (y-axis, only MA10 infection) comparisons. Arrows indicate direction of dysregulation for each comparison. Directly correlated genes (blue) occur in the same direction for both comparisons (i.e., both up-regulated or both down-regulated), while inversely correlated genes (orange) occur in opposite directions for each comparison. Linear regression line demonstrates the relationship between the two comparisons. (**B**) Venn diagram depicting the significant up- and down-regulated genes unique to the 5xFAD Control vs. WT Control (left, white) and WT MA10 vs. WT Control (right, blue), while demonstrating up- and down-regulated genes commonly shared across the two comparisons (middle).(**C**) Scatterplot of the average difference of all microglia for all significant genes between 5xFAD Control vs. WT Control, now with 5xFAD MA10 vs. 5xFAD Control (y-axis, both amyloid and MA10 infection). Red text demonstrates genes up-regulated in the same direction in the first scatterplot, but down-regulated in the MA10 5xFAD vs. Control 5xFAD comparison. (**D**) Venn diagram depicting the significant up- and down-regulated genes unique to 5xFAD Control vs. WT Control and 5xFAD MA10 vs. 5xFAD Control (right, purple), while demonstrating up-and down-regulated genes commonly shared between the two comparisons. (**E**) Three-way Venn diagram of all significantly up- and down-regulated genes within microglia between the three comparisons: 5xFAD Control vs. WT Control (white), WT MA10 vs. WT Control (blue), and 5xFAD MA10 vs. 5xFAD Control (grey). Genes are considered significantly correlated if the log2 Fold Change magnitude is greater than 0.3. (**F**) Scatterplot of the average difference of all astrocytes for all significant genes between 5xFAD Control vs. WT Control and WT MA10 vs. WT Control. (**G**) Venn diagram depicting the significant up- and down-regulated genes in astrocytes unique to 5xFAD Control vs. WT Control (left, white) and WT MA10 vs. WT Control (right, blue), while demonstrating shared dysregulated genes between the two comparisons. (**H**) Scatterplot of the average difference of all astrocytes for all significant genes between 5xFAD Control vs. WT Control. (**I**)Venn diagram depicting the significant up- and down-regulated genes in astrocytes unique to 5xFAD Control vs. WT Control (left, white) and 5xFAD MA10 vs. 5xFAD Control (right, grey), while demonstrating shared dysregulated genes between the two comparisons. (**J**) Three-way Venn diagram of all significantly up- and down-regulated genes within astrocytes between the three comparisons: 5xFAD Control vs. WT Control (white), WT MA10 vs. WT Control (blue), and 5xFAD MA10 vs. 5xFAD 48 Control (grey).

To investigate the effect of the concurrence of both inflammatory stimuli (e.g., amyloid pathology and MA10 infection) on the gene expression within microglial cells, we plotted all DEGs between Control 5xFAD vs. Control WT (only amyloid pathology) and MA10 5xFAD vs. Control 5xFAD (both amyloid pathology and MA10 infection) (**Figure 7C**). Many differential gene expression changes were shared between the two conditions, with several common genes up-regulated or down-regulated. For example, 17 genes were significantly up-regulated within myeloid cells *(Apoe, Iapp, Cast, Cblb, Neu4, Spp1, Ltbp1, Prex1, etc.)* in the presence of both inflammatory stimuli together, while 5 genes were significantly down-regulated in both *(Csf1r, P2ry12, Cx3cr1, Tgfbr1, Hexb)* (**Figure 7D**).

To determine which set of DEGs in microglial cells were unique to the interaction of both Aβ pathology and MA10 infection, we compared significant DEGs (adjusted p-value<0.05) from all microglial cells across the three comparisons for the three conditions: MA10 WT vs. Control WT (only peripheral MA10 infection), Control 5xFAD vs. Control WT (only Aβ pathology), and MA10 5xFAD vs. Control 5xFAD (both peripheral MA10 infection and Aβ pathology). As previously demonstrated, this analysis also demonstrated four homeostatic genes (*Csf1r, P2ry12, Hexb, Cx3cr1*) were significantly downregulated in the context of all three conditions. Among DEGs that were uniquely down-regulated only in the context of both amyloid pathology and MA10 infection together were *Cd74, Cst7, Tyrobp, Ctsb, H2-Aa, Cd9, C1qc, Tmsb4x,* and *Cst3.* Meanwhile, these DEGs were significantly up regulated in the presence of Aβ pathology or peripheral MA10 infection, independently (**Figure 7E**). To determine the effects on other cell types within the CNS, we performed a similar analysis with astrocytes (**Figure 7F-J**), vascular cells, oligodendrocytes, and inhibitory neurons **(Supplemental Figure 8).**

Astrocytes showed substantial overlap in transcriptional suppression across conditions. Nine down-regulated DEGs were commonly downregulated in response to either Aβ pathology or MA10 infection (*Vegfa, Atp1a2, Slc6a11, Gja1, Slc6a11, Gpr37l1, Plpp3, Gstm1, Agt*) (**Figure 7G**). MA10 infection induced ten additional down-regulated DEGs (*Sparcl1, Slc4a4, Ndrg2, Ptprz1, Slc1a2, Sox9, Aldoc, Serpine2, Cpe, Mt3)* and one upregulated DEG (*Mbp*), while Aβ pathology alone resulted in six down-regulated (*Hmgb1, Malat1, Mfge8, Bcan, Mertk, Ptn*) and ten up-regulated DEGs (*Clu, Gfap, Psen1, Tmsb4x, Vim, Apoe, Aqp4, Ctsd, Dbi, Cd81*). Six genes were commonly down-regulated with both stimuli (*Atp1a2, Mfge8, Bcan, Gja1, Gpr37l1, Plpp3*) (**Figure 7I**), and four genes (*Atp1a2, Gpr37l1, Gja1, Plpp3*) were consistently downregulated across all three conditions (**Figure 7J**).

Vascular cells shared six down-regulated DEGs in response to either Aβ pathology or MA10 infection, (*Rgs5, Slc1a2, Flt1, Sparcl1, Bsg, Pecam1*) (**Supplemental Figure 8B**). MA10 infection resulted in six additional down-regulated (*Cpe, B2m, Pltp, Sparc, Cldn5, Vtn*) and four up-regulated DEGs (*Slc2a1, Vim, Apod, Acta2*), whereas Aβ pathology alone produced one down-regulated (*Slc2a1*) and three upregulated DEGs (*Gfap, Psen1, Clu*) (**Supplemental Figure 8B**). Three genes (*Flt1, Bsg, Pecam1*) were consistently downregulated under both stimuli and across all three conditions (**Supplemental Figure 8D-E**). Oligodendrocytes showed no shared DEGs between Aβ pathology and MA10 infection (**Supplemental Figure 8G**). MA10 infection induced six down-regulated (*Pllp, Malat1, Myrf, Mag, Ptgds, Ugt8a*) and two upregulated DEGs (*Sgk1, Mbp*), while Aβ pathology resulted in one down-regulated (*Hmgb1*) and ten up-regulated DEGs (*Apoe, B2m, Clu, Cstd, Gfap, Psen1, Pllp, Cst7, Cd9, Plp1*) (**Supplemental Figure 8G**). No shared DEGs were detected under combined or all three conditions (**Supplemental Figure 8J**). Inhibitory neurons shared three down-regulated DEGs in response to either Aβ pathology or MA10 infection, (*Atp2a2, Hspa8, Sst*) **(Supplemental Figure 8L).** MA10 infection induced seven additional down-regulated (*Hsp90aa1, Tcf7l2, Gnas, Sparcl1, Ywhag, Cpe, Sparc*) and four upregulated DEGs (*Ppp1r1b, Penk, Arpp21, Pcp4*), whereas Aβ pathology resulted in five down-regulated (*Cplx1, Syp, Meg3, Snhg11, Penk*) and one upregulated DEG (*Psen1*) (**Supplemental Figure 8L**). With both stimuli, two DEGs were commonly downregulated (*Penk, Sst*) (**Supplemental Figure 8N**) and only *Sst* remained downregulated across all three conditions (**Supplemental Figure 8O**).

## DISCUSSION

As previously mentioned, recent studies have reported that infection with either SARS-CoV-2 or mouse-adapted SARS-CoV-2 (MA10) has led to the presence of gliosis and other CNS changes in infected animal models in the absence of virus in the brain at various time points arguing that peripheral inflammatory responses in response to viral infection can impact CNS resident cell response ^63–67^. While COVID-19 most commonly presents with respiratory symptoms, neurological symptoms have been extensively documented, with symptoms manifesting during the initial period of disease and lasting beyond viral clearance ^32–35,37,38,97,98^. The full extent of neuropathological changes in the CNS that results from SARS-COV-2 infection is still not fully understood. Given the association of COVID-19 with neuroinflammation and other neurodegenerative processes, it is imperative that the changes in the CNS resulting from infection and mechanisms contributing to neurodegenerative processes be more fully characterized.

In attempt to address some of these questions, we utilized a novel mouse-adapted SARS-COV-2 (MA10) to infect aged C57BL/6 (WT) mice and 5xFAD mice to examine neuropathological changes in the brain after chronic infection and effects of infection on gene expression in the brain. As both microglia and astrocytes can contribute to neuroinflammatory and neurodegenerative processes ^99–102^ changes in microglia and astrocyte number and volume were quantified. Neither microglia numbers or volume were increased within the brains of MA10 infected WT or 5xFAD mice compared to uninfected mice. Prior studies have described increases in microglia number and morphological changes suggesting that MA10 infection can induce microgliosis ^63–67^. Klein and colleagues have shown that microglia-derived IL-1β contributes to synaptic loss in the brains of infected mice ^65,66^. IL-1β is associated with impaired neurogenesis and hippocampal dependent memory ^103–105^, has been correlated with Aβ accumulation and neurofibrillary tangles (NFTs) ^106^ and can result in the aberrant release of glutamate that results in neuronal death in many neurodegenerative disease ^107^. However, in this study there were no detectable differences in IL-1β plasma levels as a result of MA10 infection. Furthermore, there was no increase in MAC2+ monocyte/macrophages within the brains of MA10-infected WT or 5xFAD mice at day 21 p.i. MA10 infection did not elicit changes in either astrocyte number or volume at day 21 p.i., and although one study has associated infection with trending increases in GFAP expression in the brains of infected laboratory mice 3 days p.i.^63^; however other reports have not found any significant increases in astrocyte numbers at 6 or 30 days p.i. ^66^. In uninfected mice, genotype does have a significant effect, leading to an increased astrocyte number and volume in the cortex of 5xFAD mice, consistent with astrocyte activation in response to increasing Aβ plaque burden ^73^. The overall lack of microglia and astrocyte activation in MA10-infected mice could be due to a variety of factors including looking at later times points rather than acute stage of disease, age of mice used for these studies, the use of a mouse-adapted variant of SARS-CoV-2 rather than naturally occurring variants that can infect mice as well as infectious dose of virus used ^108,109^.

Both MA10-infected WT and 5xFAD mice exhibited changes in gene expression that were associated with neuronal dysfunction. RNASeq analysis indicates an upregulation in genes associated with reduced long-term potentiation (LTP) in WT mice, and downregulation genes associated with synaptic function and plasticity in 5xFAD mice. LTP is a well-characterized form of synaptic plasticity and has been considered as a cellular mechanism underlying learning and memory, and LTP dysfunction is thought to be an underlying cause of memory loss ^110^. Therefore, we posit that impairments in LTP and other genes associated with synaptic plasticity were contributing factors to impaired memory noted in both human and animal studies. Similar changes in pathways associated with neuronal and synaptic dysfunction were seen in the brains of infected 5xFAD mice. The 5xFAD mice surprisingly had significantly more genes affected at day 21 p.i. in comparison to WT mice and this most likely reflects responses to Aβ plaque burden.

The RNASeq analysis of MA10 infected brains indicated neuronal and synaptic dysfunction, at least at the transcriptional level and as a result we believed these changes might also reflect changes in existing pathology. However, our findings indicate MA10 infection did not induce or affect dystrophic neurites or Aβ plaque pathology in WT or 5xFAD mice, respectively. We believed MA10 could perhaps affect these given that some coronavirus viral proteins – such as the envelope (E) protein are believed to be amyloidogenic ^24^ and animal studies indicate that MA10 infection has been reported to affect other AD-associated pathology including phosphorylated tau ^64,67^. Moreover, we did not detect increased neuronal loss nor evidence of synapse loss in infected WT and 5xFAD mice arguing that MA10 infection does not induce overt neurodegeneration at the dosage employed. Although no overt signs of increased neuropathology were detected, widespread transcriptomic dysregulation revealed more subtle molecular alterations in multiple CNS-resident cell types. Specifically, significant changes in the expression of homeostatic genes were observed across distinct cellular populations, including glial cells (microglia, astrocytes, and oligodendrocytes), inhibitory neurons, and vascular endothelial cells. These changes implicate early molecular signatures of neuroinflammation and homeostatic disruption that may represent antecedents to longer-term neurodegenerative processes. In both infected WT and 5xFAD mice, microglia exhibited downregulation of established homeostatic genes, including *Hexb*, *Csf1r*, *P2ry12*, and *Cx3cr1,* which are typically suppressed during microglial activation or a transition into a disease-associated phenotype (DAM) ^79,111,112^. *Hexb* is essential for proper lysosomal degradation, and downregulation of its expression in microglia would impair phagocytic capacity ^113^. Notably, *Hexb* encodes a myeloid-derived lysosomal enzyme which is essential for maintaining neuronal homeostasis and is not typically downregulated under pathological conditions ^114^. *Csf1r* encodes the colony-stimulating factor 1 receptor, essential for microglial survival and maintenance ^115^, while *P2ry12* and *Cx3cr1* are implicated in microglial migration and communication with neurons ^116–120^. Downregulation of these markers might suggest a shift toward a pro-inflammatory or reactive phenotype, consistent with findings in both viral encephalitis and neurodegenerative disorders such as AD ^121,122^. Furthermore, only in the MA10 infected 5xFAD brains did microglia exhibit an upregulation of genes associated with immune regulation (*Cblb*)^123^, protease inhibition (*Cast*)^124^, lysosomal degradation (*Neu4*)^125^, and inflammatory signaling/metabolic dysfunction (*Iapp*)^126,127^, indicating an additional shift away from a homeostatic state. The increased expression of *Apoe*, a hallmark of disease-associated microglia (DAM), supports the emergence of a reactive phenotype ^128^. Upregulation of *Ltbp1* and *Smurf1* suggests modulation of TGF-β signaling pathways, which play key roles in immune homeostasis ^129–133^. Elevated *Eif4g3* points to increased translational activity^134,135^ while *Ngf* upregulation may reflect a neurotrophic response^136–138^.

Astrocytes showed downregulation of key genes involved in neurotransmitter regulation (*Slc1a2*), gap junctional communication (*Gja1*), phospholipid metabolism (*Plpp3*), ion homeostasis (*Atp1a2*), and neurodevelopment (*Ptprz1*). *Slc1a2* encodes EAAT2/GLT-1, the primary glutamate transporter in astrocytes, and its downregulation has been linked to excitotoxicity and neuronal injury in several CNS diseases as well as normal aging^139–141^. Additionally, downregulation of *Atp1a2* in astrocytes impairs crucial functions like potassium clearance, leading to altered neuronal excitability ^142,143^. *Gja1*, which encodes connexin 43, is crucial for astrocyte communication and gliotransmission with neurons. Its downregulation is linked to neuroinflammation and has been shown to impair metabolic support and synaptic plasticity, contributing to neurodegenerative diseases ^81,144,145^. *Plpp3* downregulation likely disrupts lipid-mediated signaling, impairing astrocyte–neuron communication, which may lead to deficits in neurotransmission without causing overt neurodegeneration ^146^. *Ptprz1* has established roles in neurodevelopmental processes like neuronal migration and dendrite morphology, along with its association with various neurodegenerative conditions ^147–149^. Together, these transcriptomic changes indicate impairment of astrocytic roles relating to neuronal support, which could further inhibit normal neuronal function in the aftermath of viral infection.

In addition to microglia and astrocytes, we determined that oligodendroglia as well as inhibitory neurons and vascular endothelial cells in both MA10-infected WT and 5xFAD mice also displayed pronounced changes in gene expression profiles compared to uninfected mice. Oligodendrocyte transcriptomic profiles revealed downregulation of myelination-associated genes, including *Mag* ^150^, *Ugt8a*^151,152^ and *Pllp*^153,154^, as well as *Malat1*, which has neuroprotective roles^155,156^. The upregulation of *Sgk1*suggests a stress or injury response ^157–159^. These changes suggest to an environment of oligodendrocyte dysfunction or demyelination and could additionally unmask key transcriptomic changes that might contribute to the white matter abnormalities seen in post-COVID imaging studies^160,161^. Inhibitory interneurons showed reduced expression of *Ndrg4*, *Sst*, and *Npy*. *Sst* (somatostatin) and *Npy* (neuropeptide Y) which are critical neuropeptides involved in regulating neuronal excitability and synaptic transmission^87,88^. Their downregulation may indicate impaired inhibitory control, potentially contributing to disrupted cortical circuitry and excitatory-inhibitory imbalance—an emerging hallmark of COVID-19-associated cognitive and affective disturbances ^162^. Similarly, the reduction in *Ndrg4*, a gene involved in synaptic vesicle trafficking and neuronal survival ^89^, aligns with findings in AD models^90–92^ and supports the presence of early synaptic vulnerability in the post-infectious brain. Endothelial cells displayed downregulation of *Bsg*, *Cldn5* and *Pecam1*, all of which are vital for blood-brain barrier (BBB) integrity and leukocyte transmigration ^163–170^. *Cldn5* encodes a tight junction protein critical for BBB permeability, and its downregulation may reflect early BBB disruption, a finding consistent with neuropathological reports in COVID-19 patients ^62^. Conversely, upregulation of *Apod* and *Acta2* may indicate vascular remodeling suggesting vascular changes in the brain not related to BBB disruption and dysfunction but associated with remodeling of affected vessels, in a maladaptive manner, due to the stress caused by an initial insult such as inflammation and infection^171–174^.

Results from this study contribute to growing information providing insight into mechanisms by which SARS-CoV-2 infection contributes to neurologic deficits. Our results have shown there are dramatic changes in genes impacting homeostatic gene functions in microglia, astrocytes, oligodendroglia as well as neurons and vascular endothelial cells. These subtle, yet potentially important, changes in cell function can occur in the absence of viral infection of the CNS and impact neurologic functions including behavior and memory. Furthermore, MA10 infection did not impact the severity of AD-associated neuropathology in the 5xFAD model, although there were numerous changes in glial and neuron gene expression within the brains. We believe that one mechanism by which this may occur is in response to the peripheral inflammation that occurs in response to MA10 infection of the lungs leading to increased pro-inflammatory cytokines/chemokines that can access the CNS and alter resident cell function. Collectively, these findings underscore the need for further research to elucidate the molecular mechanisms underlying COVID-19-related CNS changes and their long-term impact on neurodegenerative processes.

## Acknowledgements

The authors thank Jaylen Michael Lee for assistance in performing and analyzing data from multiplex cytokine profiling.

---

**Supplemental Figure 1.**
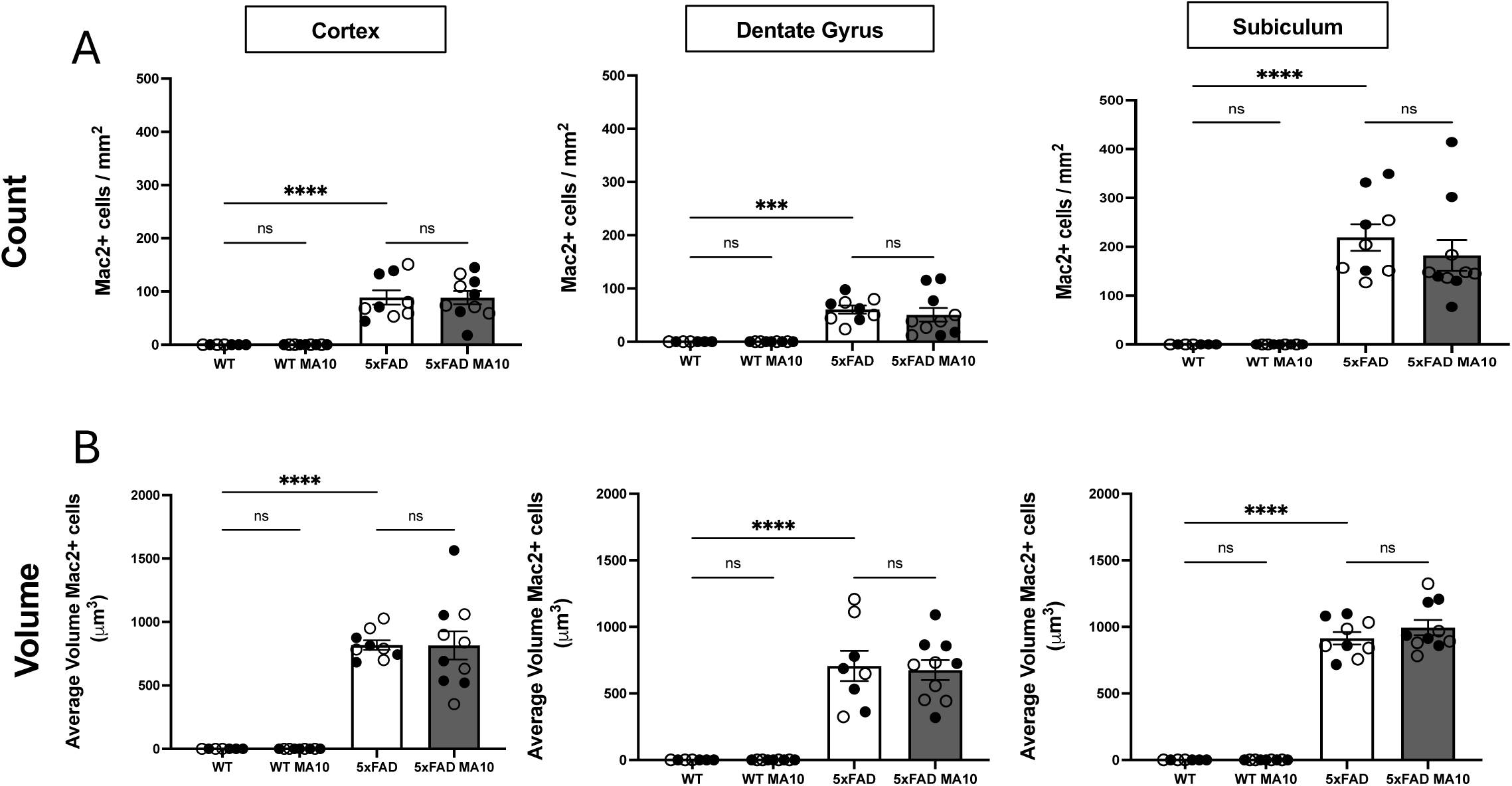
(**A**) Quantification of Mac2⁺ cells and their volumes (**B**) in the cortex, dentate gyrus, and subiculum of MA10-infected WT and 5xFAD mice at day 21 p.i. Immunohistological data were analyzed using two-way ANOVA. Tukey’s post-hoc test was employed to examine biologically relevant interactions. Female and male mice are indicated by open or closed circles respectively. Data are represented as mean ±SEM; *** p≤0.001, **** p≤0.0001.

**Supplemental Figure 2.**
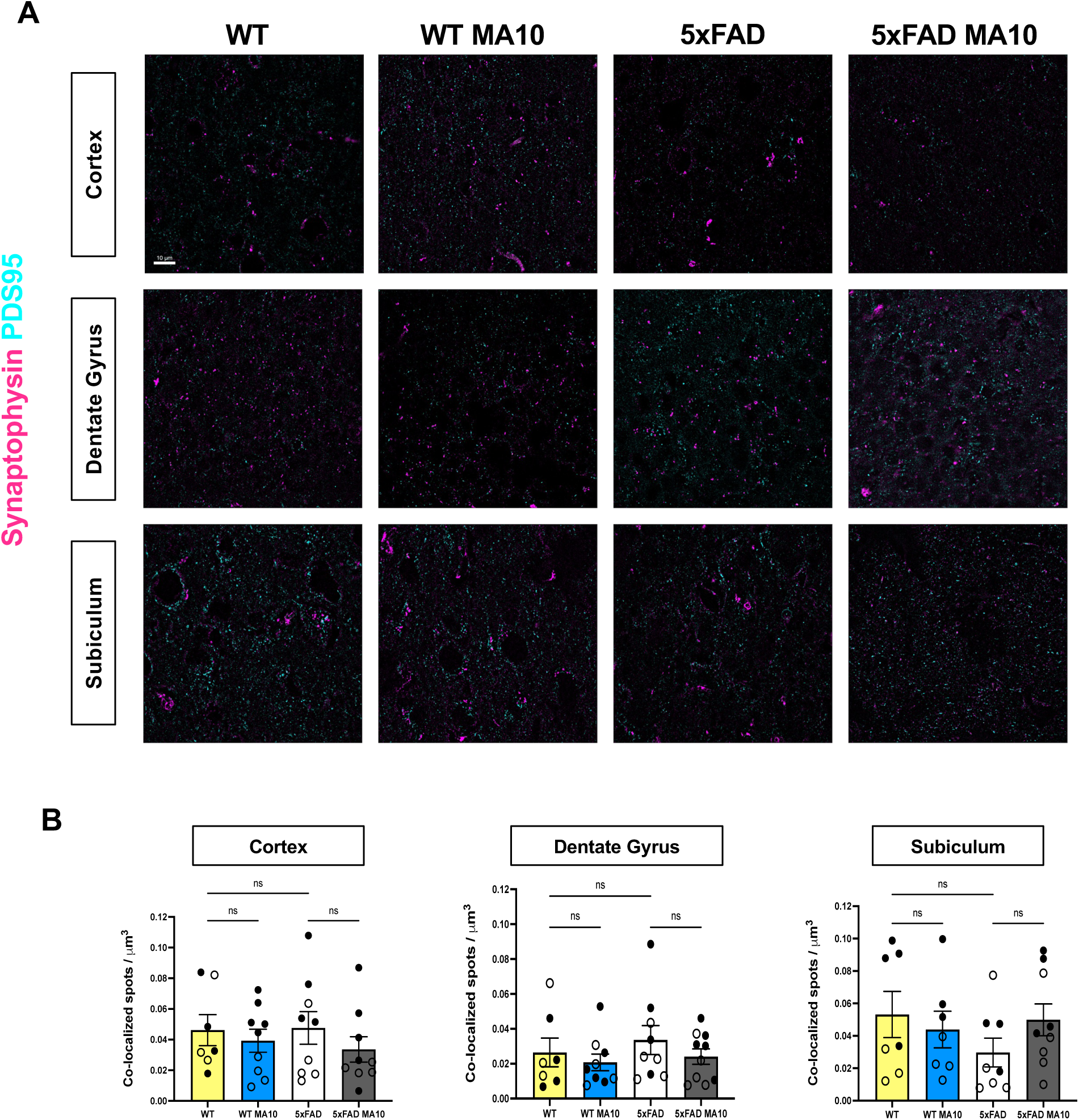
MA10 infection and synaptic vesicle density in WT and 5xFAD mice. Brains of MA10-infected uninfected WT and 5xFAD mice at day 21 p.i. were immunostained with synaptophysin for presynaptic elements (magenta) and PSD-95 for postsynaptic elements (turquoise). (**A**) Representative super-resolution images at 40X objective of cortex, dentate gyrus, and subiculum from infected and uninfected mice are shown. (**B**) Quantification of synaptophysin+ and PSD-95+ colocalized spots per µm3. Immunohistological data were analyzed using two-way ANOVA. Tukey’s post-hoc test was employed to examine biologically relevant interactions. Female and male mice are indicated by open or closed circles respectively. Data are represented as mean ±SEM. Scale bar in (A) = 10 µm.

**Supplemental Figure 3.**
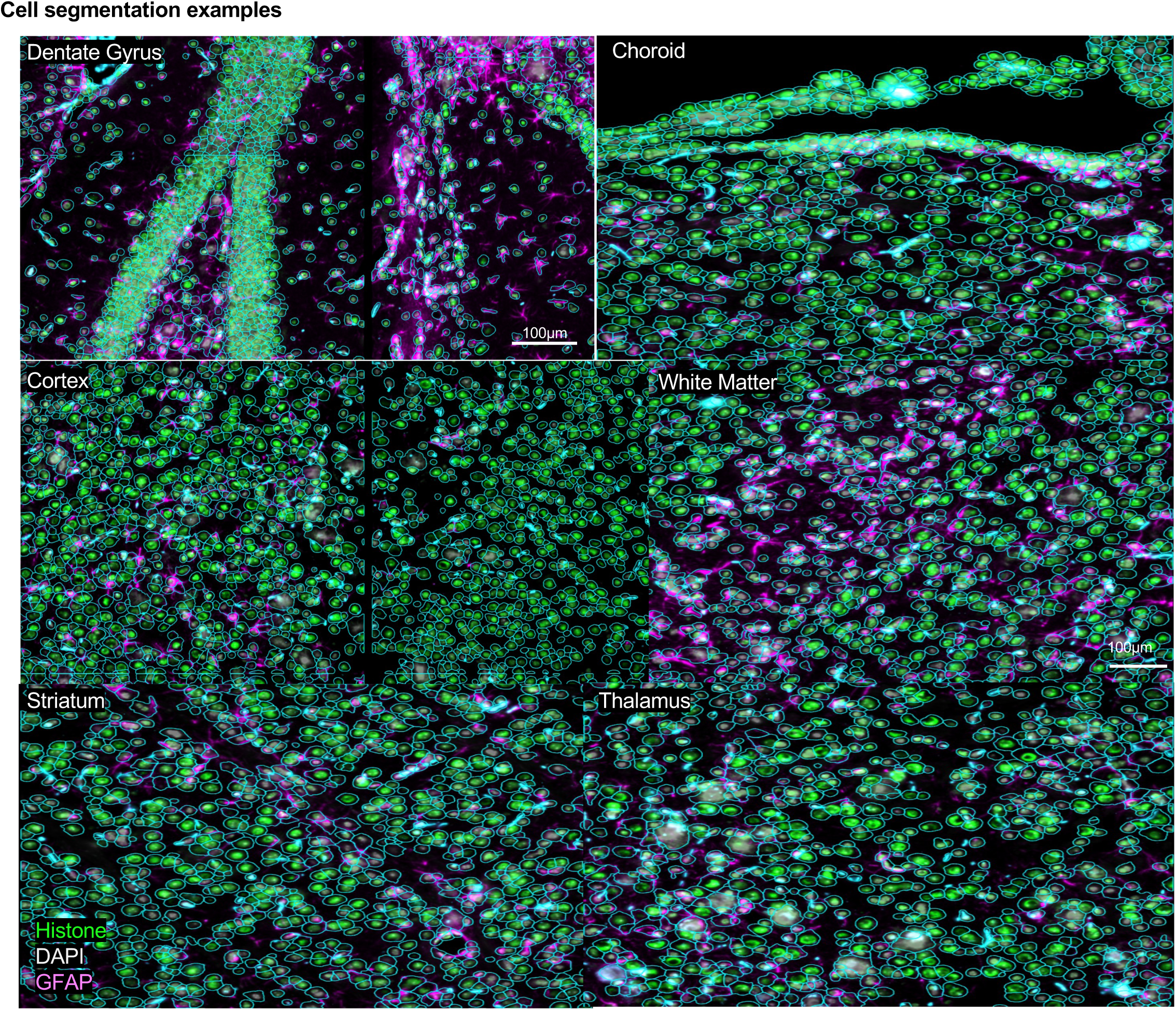
Representative images demonstrating cell segmentation in dentate gyrus, choroid plexus, cortex, white matter, striatum and thalamus. Cells were imaged with rRNA (not shown), histone, DAPI, and GFAP markers and segmented automatically.

**Supplemental Figure 4.**
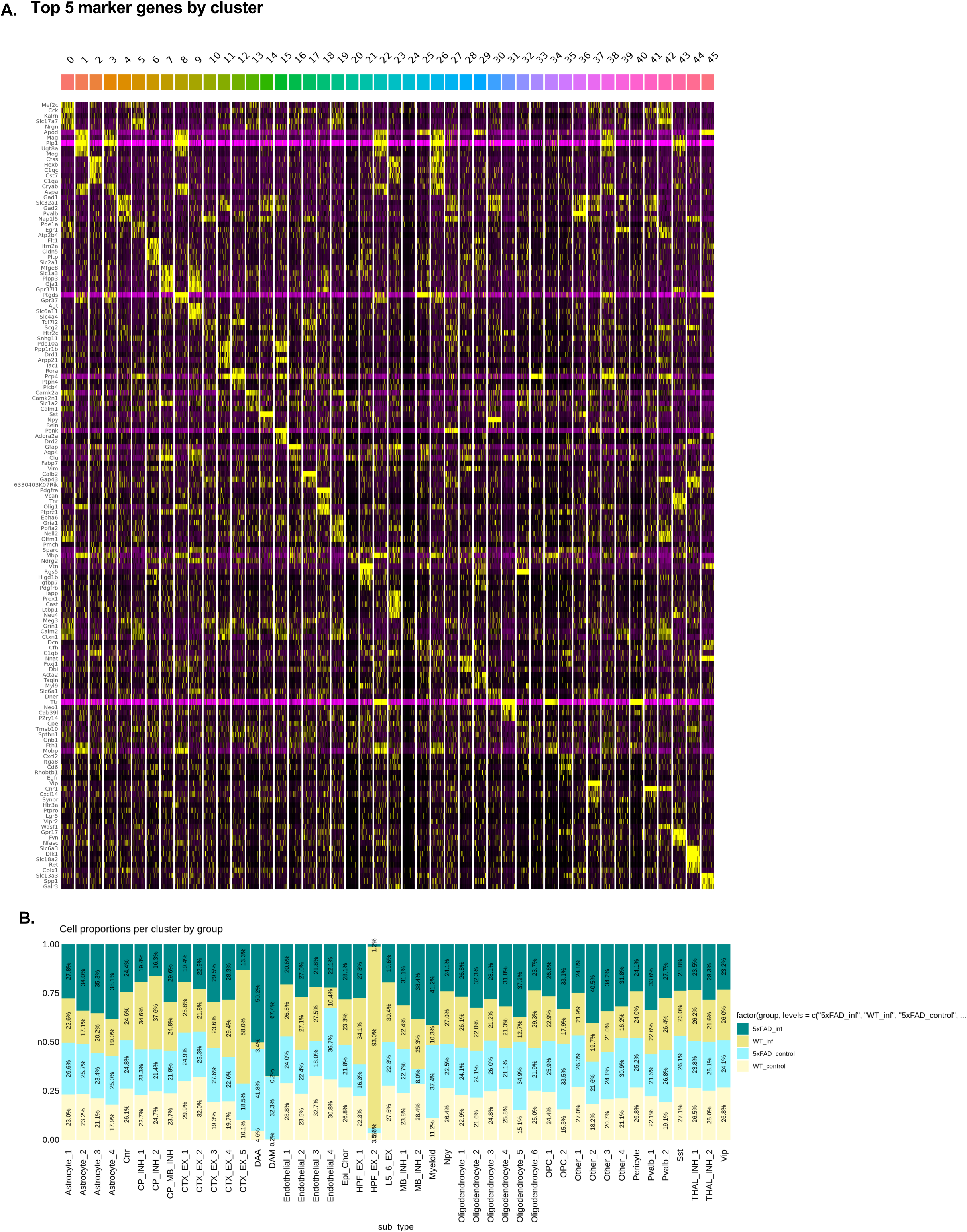
(**A**) Heatmap of top 5 marker genes for all spatial transcriptomics subclusters. (**B**) Proportional distribution of Seurat clusters across experimental groups. Bar plots display relative abundance of each cluster in each group.

**Supplemental Figure 5.**
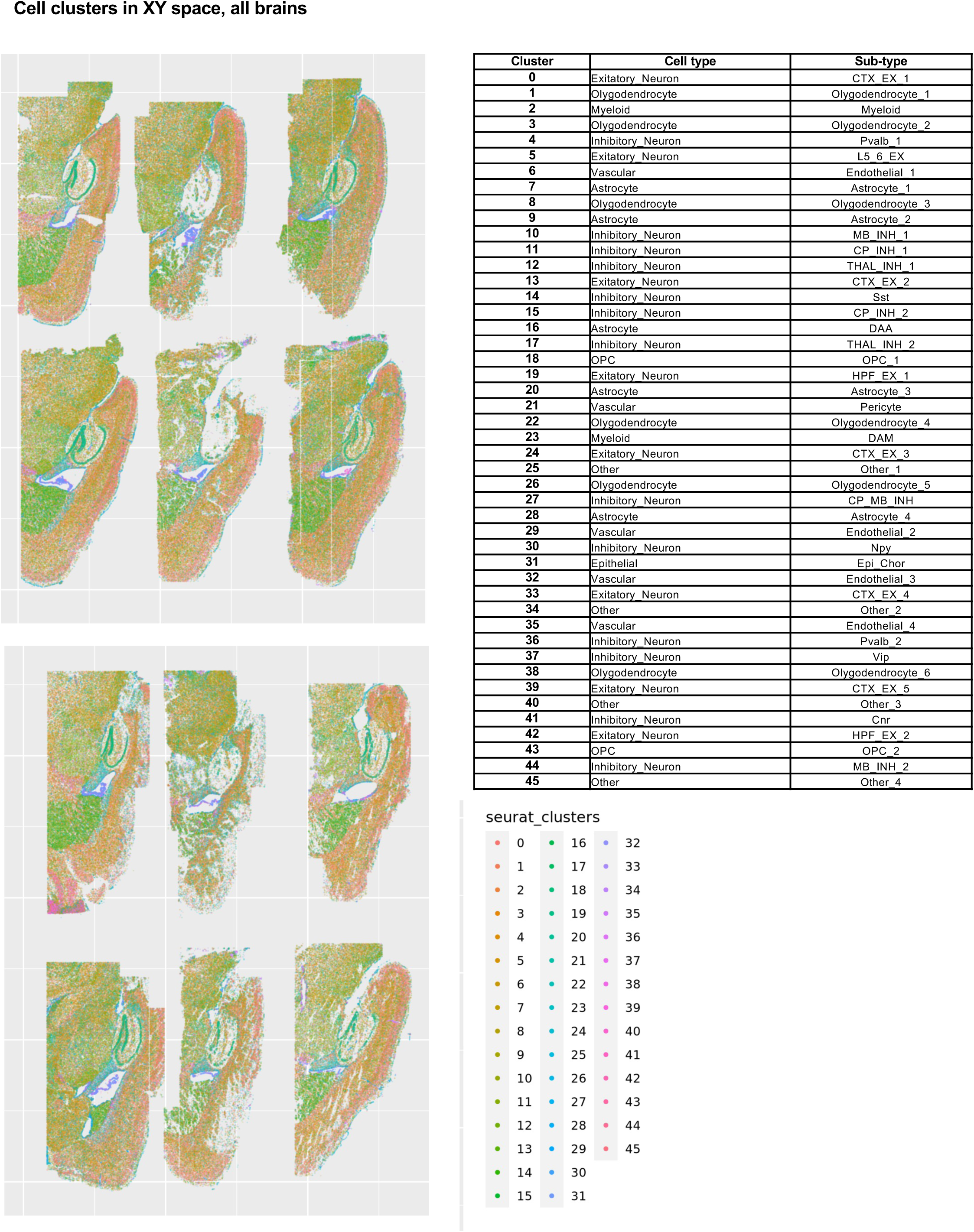
46 clusters plotted in XY space in all 6 brains from WT (n=3) and 5xFAD (n=3) mice, both infected and uninfected, at day 21 p.i. (n=3/group).

**Supplemental Figure 6.**
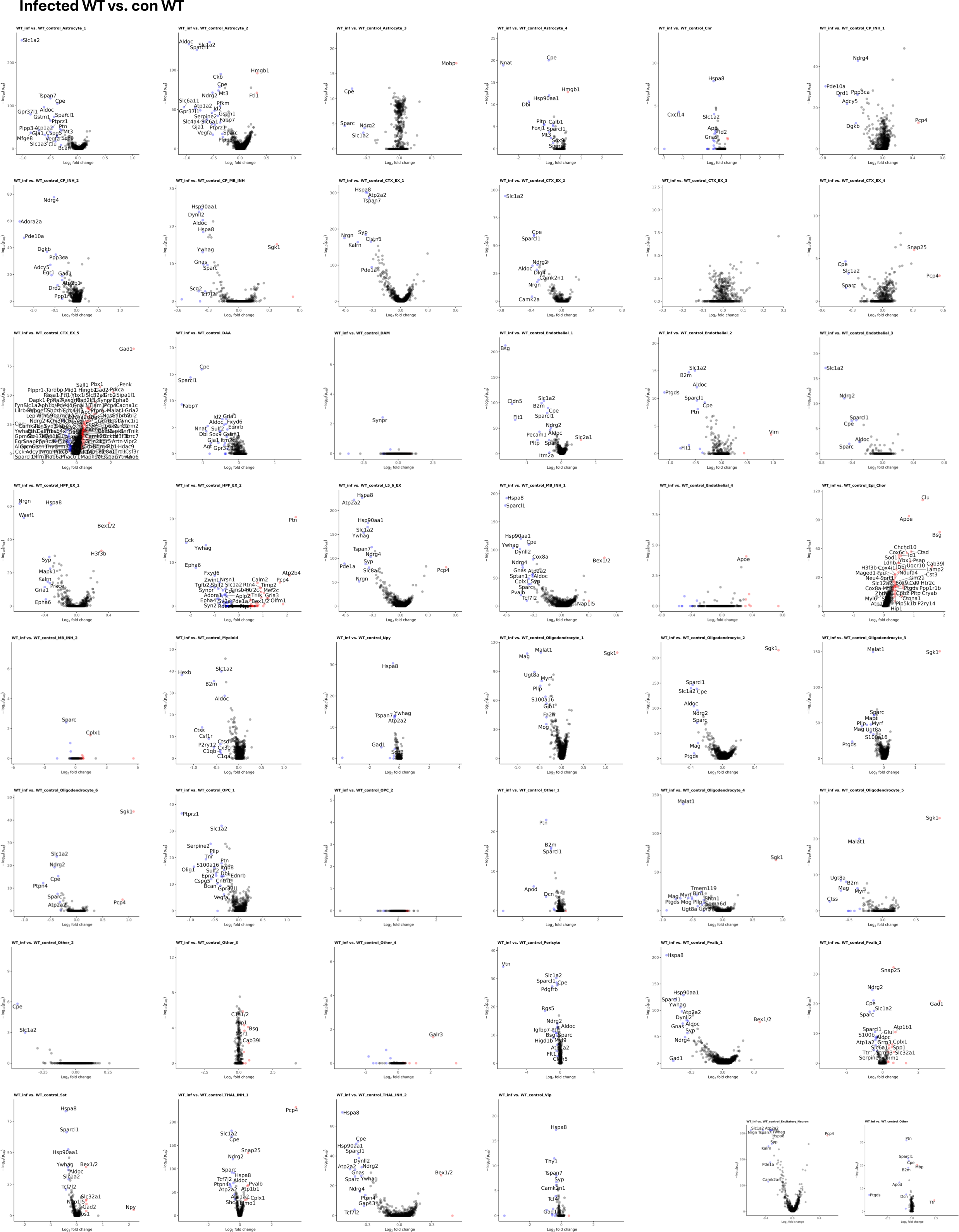
Volcano plots for all spatial transcriptomics subcluster in MA10 infected WT vs Control WT.

**Supplemental Figure 7.**
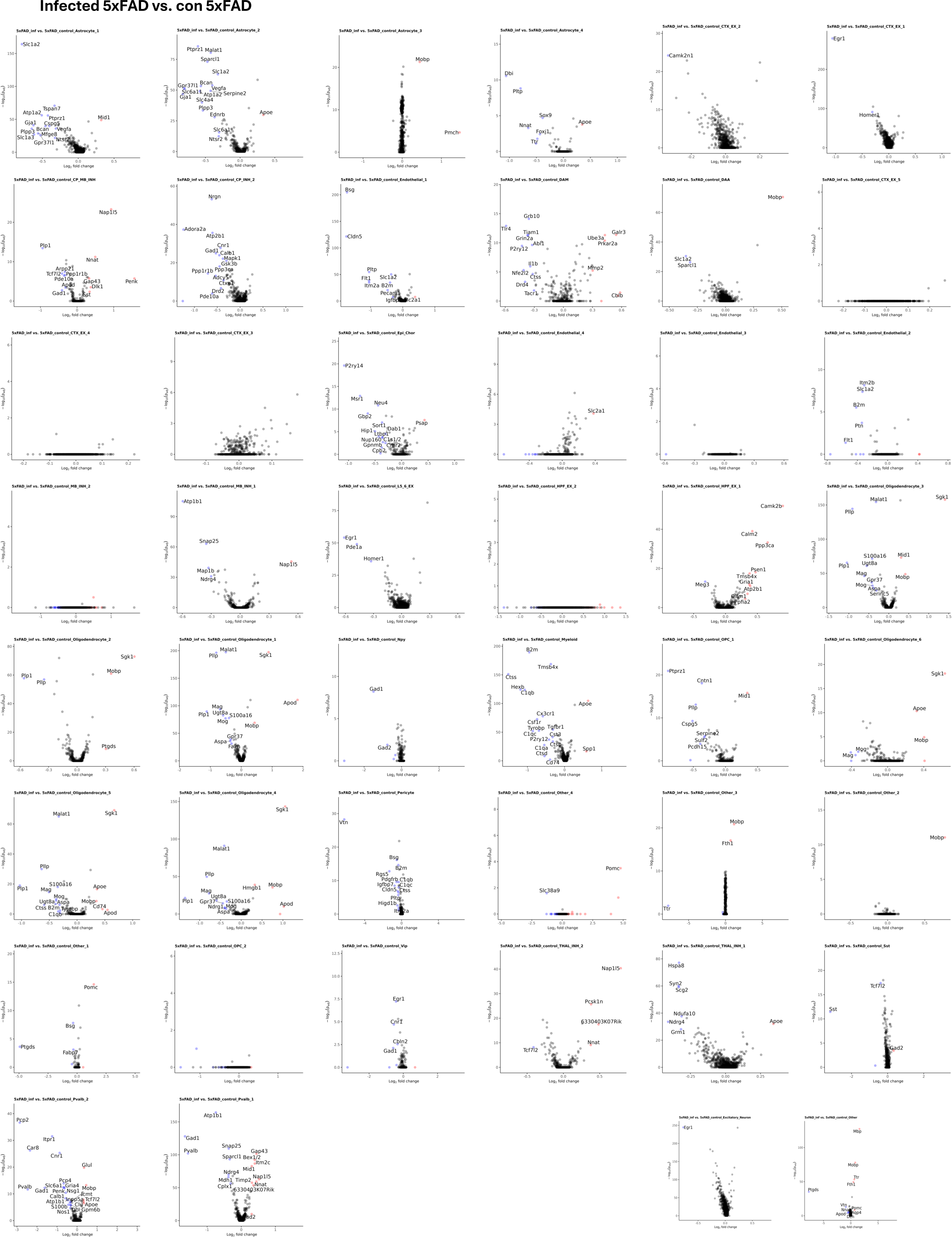
Volcano plots for all spatial transcriptomics subcluster in MA10 infected 5xFAD vs Control 5xFAD.

**Supplemental Figure 8.**
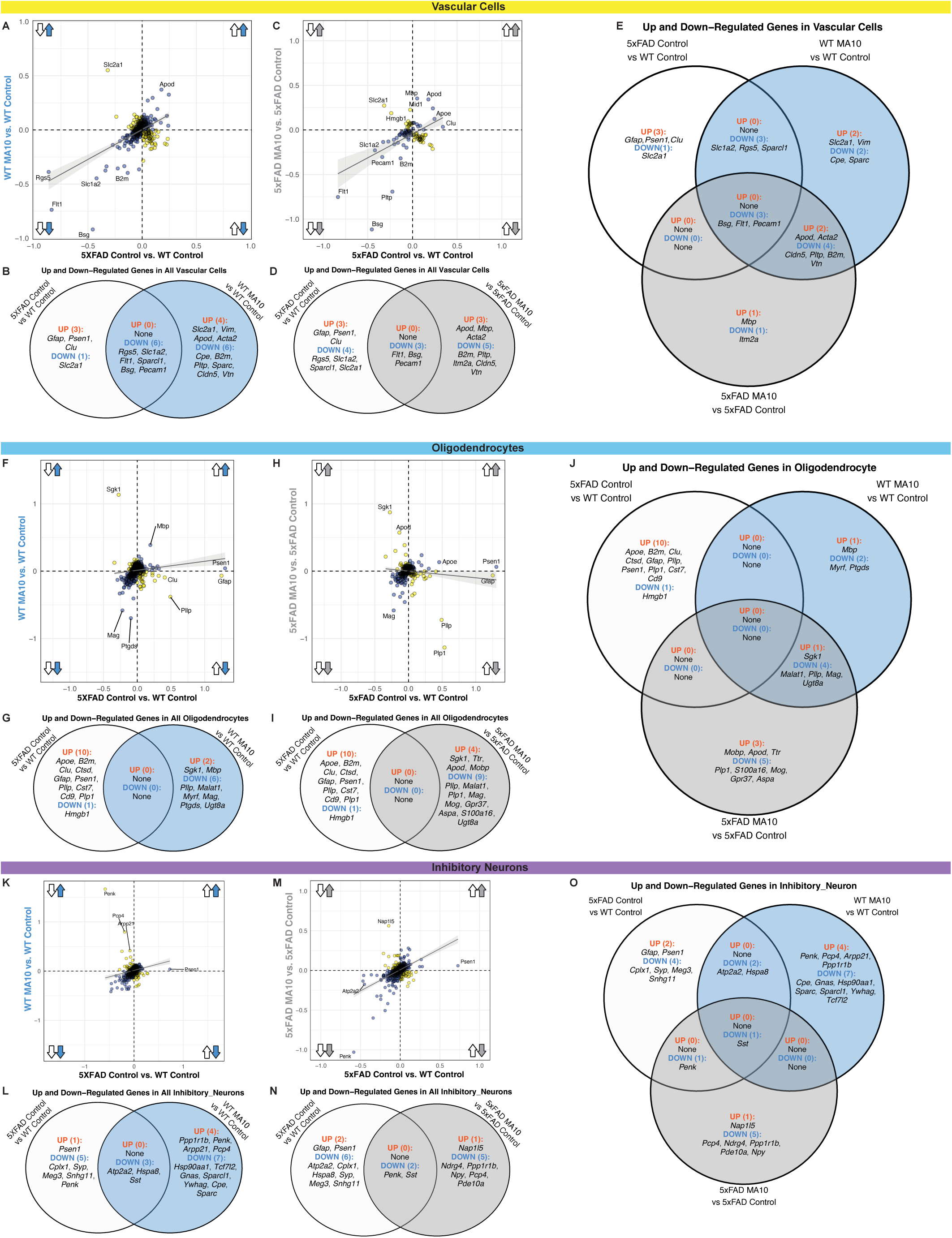
Investigating gene dysregulation within vascular cells, oligodendrocytes and inhibitory neurons in the presence of amyloid pathology, peripheral MA10 infection, or both concurrently. (**A**) Scatterplot of the average difference of all vascular cells for all significant genes between 5xFAD Control vs. WT Control and WT MA10 vs. WT Control. (**B**) Venn diagram depicting the significant up-and down-regulated genes in vascular cells unique to 5xFAD Control vs. WT Control (left, white) and WT MA10 vs. WT Control (right, blue), while demonstrating shared dysregulated genes between the two comparisons. (**C**) Scatterplot of the average difference of all vascular cells for all significant genes between 5xFAD Control vs. WT Control. (**D**) Venn diagram depicting the significant up-and down-regulated genes in vascular cells unique to 5xFAD Control vs. WT Control (left, white) and 5xFAD MA10 vs. 5xFAD Control (right, grey), while demonstrating shared dysregulated genes between the two comparisons. (**E**) Three-way Venn diagram of all significantly up- and down-regulated genes within vascular cells between the three comparisons: 5xFAD Control vs. WT Control (white), WT MA10 vs. WT Control (blue), and 5xFAD MA10 vs. 5xFAD Control (grey). (**F**) Scatterplot of the average difference of all oligodendrocytes for all 50 significant genes (padj < 0.05) between 5xFAD Control vs. WT Control (x-axis, only amyloid pathology) and WT MA10 vs. Control WT (y-axis, only MA10 infection) comparisons. Arrows indicate direction of dysregulation for each comparison. Directly correlated genes (blue) occur in the same direction for both comparisons (i.e., both up-regulated or both down-regulated), while inversely correlated genes (orange) occur in opposite directions for each comparison. Linear regression line demonstrates the relationship between the two comparisons. (**G**) Venn diagram depicting the significant up- and down-regulated genes unique to the 5xFAD Control vs. WT Control (left, white) and WT MA10 vs. WT Control (right, blue), while demonstrating up- and down-regulated genes commonly shared across the two comparisons (middle). (**H**) Scatterplot of the average difference of all oligodendrocytes for all significant genes between 5xFAD Control vs. WT Control, now with 5xFAD MA10 vs. 5xFAD Control (y-axis, both amyloid and MA10 infection). Red text demonstrates genes up-regulated in the same direction in the first scatterplot, but down-regulated in the MA10 5xFAD vs. Control 5xFAD comparison. (**I**) Venn diagram depicting the significant up- and down-regulated genes unique to 5xFAD Control vs. WT Control and 5xFAD MA10 vs. 5xFAD Control (right, purple), while demonstrating up- and down-regulated genes commonly shared between the two comparisons. (**J**) Three-way Venn diagram of all significantly up- and down-regulated genes within oligodendrocytes between the three comparisons: 5xFAD Control vs. WT Control (white), WT MA10 vs. WT Control (blue), and 5xFAD MA10 vs. 5xFAD Control (grey). Genes are considered significantly correlated if the log2 Fold Change magnitude is greater than 0.3.(**K**) Scatterplot of the average difference of all inhibitory neurons for all significant genes between 5xFAD Control vs. WT Control and WT MA10 vs. WT Control. (**L**) Venn diagram depicting the significant up- and down-regulated genes in inhibitory neurons unique to 5xFAD Control vs. WT Control (left, white) and WT MA10 vs. WT Control (right, blue), while demonstrating shared dysregulated genes between the two comparisons. (**M**) Scatterplot of the average difference of all inhibitory neurons for all significant genes between 5xFAD Control vs. WT Control. (**N**) Venn diagram depicting the significant up- and down-regulated genes in inhibitory neurons unique to 5xFAD Control vs. WT Control (left, white) and 5xFAD MA10 vs. 5xFAD Control (right, grey), while demonstrating shared dysregulated genes between the two comparisons. (**O**) Three-way Venn diagram of all significantly up- and down-regulated genes within inhibitory neurons between the three comparisons: 5xFAD Control vs. WT Control (white), WT MA10 vs. WT Control (blue), and 5xFAD MA10 vs. 5xFAD Control (grey).

